# Nanochirality-programmed Type-I photosensitizer enables deep-tumor photodynamic therapy by reducing extracellular-matrix adhesion

**DOI:** 10.1101/2025.10.01.679807

**Authors:** Yichen Liu, Gaeun Kim, Runyao Zhu, Hyunsu Jeon, Yichun Wang

## Abstract

Tumor hypoxia and poor penetration of therapeutics across tumor-microenvironment barriers remain major obstacles to effective cancer therapy, including photodynamic therapy (PDT). Here we introduce a nanochirality-programmed assembly (*L*-Chi-GAIN) in which nanochirality drives site-selective assembly that activates oxygen-independent Type-I reactive oxygen species (ROS) generation and reduces hyaluronan-mediated matrix adhesion, thereby permitting deep intratumoral therapy. Glycosylation imparts structural chirality to graphene quantum dots (GQDs), directing site-selective assembly of indocyanine green (ICG) that turns on photoinduced electron transfer (PET), producing a 64-fold increase in ROS relative to free ICG. Nanochirality also modulates assembly–extracellular matrix (ECM) interactions. *L*-GQDs show a less favorable hyaluronan binding free energy (ΔG_bind_), thus accelerating interstitial transport and resulting in ∼21-fold deeper tumor penetration by *L*-Chi-GAIN than conventional nanocarriers. Under near-infrared irradiation, *L*-Chi-GAIN elicits strong oxidative stress and triggers Gasdermin-D (GSDMD)-dependent pyroptosis, leading to significant suppression of tumor growth. This work offers a nanochirality-guided design strategy for PDT in deep tumors by coupling site-selective assembly with stereoselective navigation of the ECM.

## 1. Introduction

Photodynamic therapy (PDT) has emerged as a clinically validated and rapidly evolving cancer treatment strategy, which offers a unique combination of advantages, including noninvasiveness, spatiotemporal precision, repeated dosing, and minimal toxicity.^1-5^ By leveraging light-activated photosensitizers (PSs) to generate cytotoxic reactive oxygen species (ROS), PDT enables localized tumor ablation and spares surrounding healthy tissues. This approach is effective for superficial malignancies and is increasingly explored for solid tumors, such as esophageal, lung, and prostate cancers.^6^ However, most clinically available PSs function via a Type-II mechanism, which depends on O_2_ to produce singlet oxygen (^1^O_2_).^7, 8^ This O_2_ dependency renders them ineffective under the hypoxic conditions commonly found in solid tumors, which limits their therapeutic efficacy.^9, 10^ Moreover, the dense extracellular matrix (ECM) and elevated interstitial fluid pressure in tumors present additional barriers, restricting the penetration and uniform distribution of PSs in deep tumor regions.^11-13^.

By comparison, Type-I PDT offers a promising strategy for overcoming hypoxic conditions in tumor microenvironments. Unlike Type-II PDT, Type-I PDT relies on electron transfer to generate superoxide anion radicals (O_2_^•–^), which dismutate into secondary ROS for cytotoxic efficacy under low-oxygen conditions and potentially contributing to local O_2_ restoration.^14-19^ Recent efforts to develop Type-I PSs have demonstrated the potential of intermolecular excitonic coupling within supramolecular dye assemblies to increase the intersystem crossing (ISC) efficiency, thereby promoting Type-I ROS generation.^20-22^ This approach minimizes structural perturbation and reduces the risk of undesired toxicity associated with the altered photochemical behavior. However, previous studies have mainly consisted of early-stage mechanistic investigations, and they have not included design considerations for tumor targeting, tissue penetration, and physiological stability, which are essential for effective PDT in solid tumors.^23^ Therefore, a new design strategy is required to afford tumor specificity, deep-tissue penetration, and optimized excited-state dynamics.

Nanotechnology offers versatile strategies to enhance cancer treatment by enabling precise control over material properties and biological interactions.^24-27^ Nanostructures with tunable size, surface chemistry, and ligand functionalization can improve pharmacokinetics and enable receptor-mediated targeting.^24-27^ Furthermore, well-defined nanoscale assemblies that organize molecules or excitons strengthen electronic coupling, prolong triplet lifetimes, thereby boosting reactive-species generation and enhancing PDT efficacy.^28, 29^ Graphene quantum dots (GQDs) have emerged as a promising nanoplatform for phototherapy owing to their high surface area, excellent biocompatibility, and tunable band structure.^30-32^ Their edge-localized π-domains and oxygenated defect sites introduce low-energy acceptor states and strong transition-dipole coupling, which facilitate efficient electron transfer.^33, 34^ Moreover, GQDs can be imparted with chirality through edge functionalization with chiral molecules, which induces nanoscale asymmetry that enables stereospecific interactions with lipid bilayers and ECM components.^35-37^ Therefore, rationally designed nanoscale chiral GQDs can achieve enhanced transmembrane transport, improved intratumoral diffusion, and higher overall delivery efficiency.

Here, we present a chiral nanomaterial composed of chiral GQDs assembled with indocyanine green (ICG) to form a nanocomplex, termed chiral graphene quantum dot–indocyanine green nanoassembly (Chi-GAIN) (**Fig. 1a**). This platform addresses the key limitations of PDT in solid tumor treatment by enabling directional electron transfer, enhancing deep intratumoral penetration, and inducing immunogenic cell death (**Fig. 1a and 1b**). ICG is an FDA-approved Cy7 dye that has been used as Type-II PS to treat hepatocellular carcinoma (HCC).^38^ However, its O_2_ dependency and low ISC efficiency limit ROS generation and reduce the PDT efficacy.^39^ To overcome this limitation, we first synthesized chiral GQDs by glycosylation, inducing out-of-plane distortion and confers nanoscale chirality. These galactose groups simultaneously act as ligands for HCC and endow nanostructures with receptor-mediated targeting capabilities. Nanoscale chirality drives the localized self-assemblyof ICG on chiral GQDs, strengthening excitonic coupling to accelerate O_2_^•–^ generation and drive light-activated NADH oxidation. The resulting *L*-Chi-GAIN nanoassembly achieves a 64-fold increase in ROS output compared to free ICG and a maximal turnover frequency of 49.8 min^−1^ for NADH oxidation, 2–3 orders of magnitude higher than commonly studied transition-metal catalysts. Equally important, *L*-GQDs form weaker adhesive contacts and exhibit shallower energy minima with hyaluronan (HA), reducing stromal retention and enhancing interstitial transport within the tumor microenvironment. This translates into ∼21-fold deeper tumor penetration relative to liposomal carriers. Moreover, *L*-Chi-GAIN triggers pyroptosis, thereby activating antitumor immunity. Together, these findings establish a generalizable design paradigm in which nanoscale chirality simultaneously modulates photochemical reactivity and tumor-ECM interactions, thereby overcoming the barriers of hypoxia and diffusion that limit conventional PDT and opening a promising path toward next-generation cancer treatment.

**Fig. 1.**
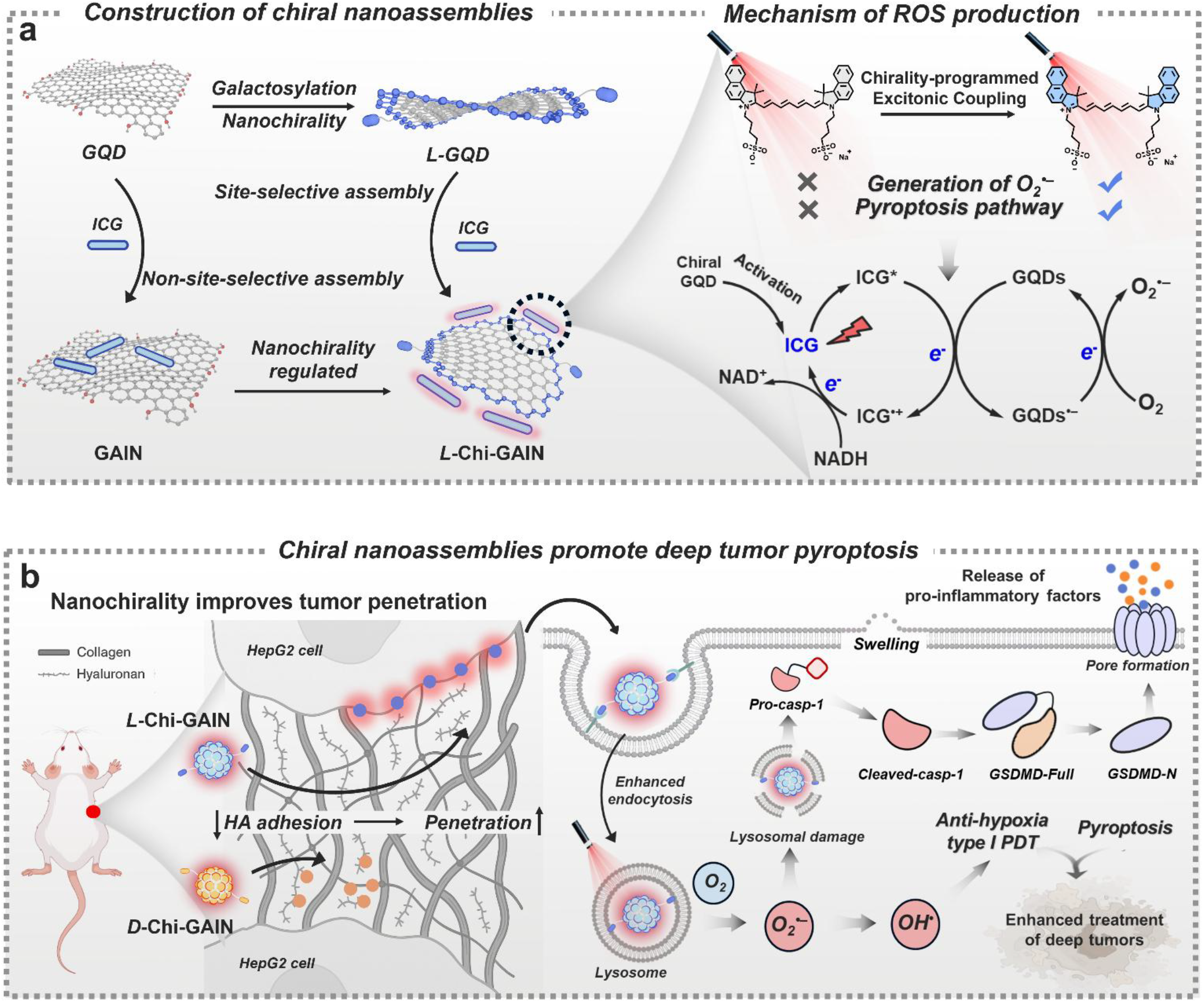
Schematic representation of photodynamic therapy (PDT) based on the chiral nano-assembly platform. (a) Chiral GQD synthesis and Chi-GAIN construction with reprogramming of the photophysics of ICG and acceleration of O_2_^•–^ generation. (b) Mechanism by which *L*-Chi-GAIN penetrates tumors and induces deep intratumoral pyroptosis to enhance PDT.

## 2. Materials and methods

### 2.1. Ethical statement

All the animal experiments were performed according to the protocols evaluated and approved by the Institutional Animal Care and Use Committee of the University of Notre Dame. (Approval Number: 24-11-8960)

### 2.2. Materials and reagents

All the reagents and solvents were obtained from commercial sources and used as received unless otherwise noted. Carbon nanofibers, nitric acid, sulfuric acid, EDC·HCl, N-hydroxysulfosuccinimide sodium salt (sulfo-NHS), HPF, 4′,6-diamidino-2-phenylindole (DAPI), and arginine were purchased from Sigma-Aldrich (St. Louis, MO, USA). Amino-L-galactose, amino-D-galactose, NADH, ABDA, DCFH-DA, AO, DPPC, DSPE-PEG_2000_, and ICG were obtained from Ambeed (Buffalo Grove, IL, USA). DHE was obtained from AAT BioQuest (Pleasanton, CA, USA). Hoechst 33342, LysoView™ 594, and DiD (DiIC_18_) were from Biotium (Fremont, CA, USA). The CCK-8 assay kit was purchased from Enzo Life Sciences (Farmingdale, NY, USA). The LIVE/DEAD Cell Imaging Kit (488/570; R37601) was purchased from Thermo Fisher Scientific (Waltham, MA, USA). Primary antibodies were purchased from Abcam (Cambridge, UK), and HRP-linked secondary antibodies were purchased from Santa Cruz Biotechnology (Dallas, TX, USA). FBS was obtained from VWR (Radnor, PA, USA). Dulbecco’s Modified Eagle Medium/Nutrient Mixture F-12 (DMEM/F-12) was purchased from Thermo Fisher Scientific, and PBS was obtained from Corning (Corning, NY, USA). Human hepatocellular carcinoma HepG2 cells (ATCC HB-8065) were obtained from the ATCC (Manassas, VA, USA).

### 2.2. In vitro O_2_^•–^ detection by DHE

PBS (2.0 mL) containing Chi-GAIN or free ICG (20 µM), DHE (40 µM), and calf-thymus DNA (ctDNA, 500 µg mL^− 1^) was transferred to a quartz cuvette. Samples were irradiated with a 730 nm light-emitting diode (LED; 20 mW cm^−2^) for the indicated times. Matched dark controls were shielded from light. The oxidation of DHE to ethidium was monitored by fluorescence (λex 560 nm; emission collected at 610 nm). Under these assay conditions, DHE oxidation predominantly reflects O_2_ ^•–^ production.

### 2.4. In vitro ^1^O_2_ detection by ABDA

ABDA (50 µM) was added to PBS solutions (2.0 mL) containing Chi-GAIN or free ICG (20 µM) in quartz cuvettes. Samples were irradiated with a 730 nm LED (20 mW cm^−2^) for the indicated times. Parallel dark controls were shielded from light. Photooxidation of ABDA by ^1^O_2_ was quantified by the decrease in absorbance at 378 nm (A_378_) recorded using a UV–Vis spectrophotometer.

### 2.5. In vitro O_2_^•–^ detection by ESR spectroscopy

DMPO (25 mM final) was used as a spin-trap for photogenerated O_2_^•–^. Aqueous samples containing 100 µM PSs and 25 mM DMPO were prepared under the following conditions: (i) *L*-Chi-GAIN + light, (ii) *D*-Chi-GAIN + light, (iii) free ICG + light, (iv) *L*-Chi-GAIN dark control, (v) *D*-Chi-GAIN dark control, and (vi) DMPO only + light control. The samples exposed to light were irradiated with a 730 nm LED (20 mW cm^−2^) for the indicated times. Immediately after treatment, the ESR spectra were recorded, and the presence of the characteristic DMPO radical adduct multiplet was taken as evidence of O_2_^•–^ generation.

### 2.6. Computational details

Geometry optimization and frontier molecular orbital (HOMO/LUMO) wavefunction analyses were performed using DFT at the B3LYP/6-31G(d) level, including Grimme’s D3 dispersion correction with the original damping function.^40^ Vertical excited-state energies were obtained using time-dependent DFT (TD-DFT) at the same level. All the electronic structure calculations were conducted using Gaussian 16.^41^ RDG surfaces and electron–hole distributions were generated using Multiwfn v3.8.^42^ All-atom molecular dynamics (MD) simulations were performed using GROMACS 2022.1 with the CHARMM36m force field for proteins, carbohydrates, and lipids, and the TIP3P water model. Topologies for *L*-GQD and *D*-GQD were generated with the CHARMM General Force Field (CGenFF). The extracellular matrix (ECM) model was constructed by assembling a collagen triple helix (Gly-Pro-Hyp repeats) and a hyaluronan (HA) polymer chain (degree of polymerization ∼50), reflecting the composition and density reported for tumor stroma. The system was solvated in a rectangular water box with 0.15 M NaCl to mimic physiological ionic strength, neutralized, and energy-minimized using the steepest-descent algorithm. All visualizations were generated using VMD 1.9.4 and PyMOL, and free-energy profiles were derived via gmx energy and reweighted using the Weighted Histogram Analysis Method (WHAM). Interaction energy profiles and contact probabilities were averaged over the final 300 ns of equilibrated trajectories.

### 2.7. In vitro cytotoxicity assay

HepG2 cells were seeded in 96-well plates at 5 × 10^3^ cells per well and allowed to attach for 24 h under normoxic (21% O_2_) or hypoxic (2% O_2_) conditions. The culture medium was then replaced with 100 µL of DMEM containing the indicated concentrations of Chi-GAIN, and the cells were incubated for 6 h. Subsequently, the wells were washed three times with PBS, replenished with fresh DMEM, and irradiated with a 730 nm LED (20 mW cm^−2^) for 10 min. After an additional 12 h of incubation, the cell viability was determined using the CCK-8 assay according to the manufacturer’s protocol. The dark toxicity controls were processed in the same way and shielded from light during the irradiation step.

### 2.8. Intracellular ROS detection

HepG2 cells were seeded on 35 mm glass-bottom confocal dishes (1 × 10^5^ cells per dish) and cultured for 24 h under normoxic (21% O_2_) or hypoxic (2% O_2_) conditions. The cells were then incubated with Chi-GAIN (20 µM in complete medium) for 4 h, washed three times with PBS, and loaded with the indicated ROS probe (general ROS: DCFH-DA (2 µM, 30 min, 37 °C, dark); superoxide: DHE (2 µM, 30 min, 37 °C, dark); and hydroxyl radical: HPF (2 µM, 30 min, 37 °C, dark)). After probe loading, the cells were washed three times with PBS and irradiated with a 730 nm LED (20 mW cm^−2^, 10 min) or kept in the dark (control). Fluorescence images were acquired immediately using CLSM with the same acquisition settings within each probe set.

### 2.9. Live/dead cell staining assay

HepG2 cells were seeded on 35 mm glass-bottom confocal dishes (1 × 10^5^ cells per dish) and cultured for 24 h under normoxic (21% O_2_) or hypoxic (2% O_2_) conditions. The cells were then incubated with Chi-GAIN (20 µM in complete medium) for 4 h and washed three times with PBS. Cultures were irradiated with a 730 nm LED (20 mW cm^− 2^, 20 min) or kept in the dark (control), and incubated for 12 h post-treatment. Live/dead staining was performed with calcein AM and PI (30 min, 37 °C, dark), the cells were washed with PBS, and fluorescence images were acquired immediately using CLSM.

### 2.10. Intracellular NADH determination

Experiments to determine the changes in NADH in the HepG2 cells treated with different concentrations of Chi-GAIN were performed using the NAD^+^/NADH assay kit with WST-8. Briefly, HepG2 cells were seeded in six-well plates, incubated for 24 h, and then exposed to Chi-GAIN in DMEM. After incubation for another 4 h, the cells were washed with PBS buffer three times, infused with fresh DMEM, and illuminated by LED light (730 nm, 20 mW cm^−2^) for 10 min. After further incubation for 4 h, the HepG2 cells were collected for subsequent NAD^+^/NADH detection using an NAD^+^/NADH assay kit. For NAD^+^/NADH extraction, 200 μL of NAD^+^/NADH extraction buffer was added, blown gently, and then spun at 12,000 rpm (5 min, 4 °C) to obtain the supernatant, which was used as the test sample. The supernatant was heated at 60 °C for 30 min, added to 96-well plates that contained the detection reagent, and incubated at 37 °C. Incubation continued until orange formazan appeared, at which point the NADH levels were quantified by measuring the absorbance at 450 nm using a microplate reader.

### 2.11. Western blot

The pyroptosis mechanism was confirmed by detecting its pathway markers (Pro-Caspase-1 and GSDMD) by western blot analysis. Briefly, proteins from the cell lysates were separated by sodium dodecyl sulfate-polyacrylamide gel electrophoresis (SDS-PAGE). For the Caspase-1 and GSDMD pathways, each well in the SDS gels was loaded with 8 and 15 µg of proteins from the cell lysates, respectively. The separated proteins were transferred onto nitrocellulose membranes. The membranes were then incubated overnight with primary antibodies, followed by incubation with horseradish peroxidase-conjugated secondary antibodies. Immunoreactive species were visualized using an enhanced chemiluminescence substrate, and bands were detected using a C600 Bioanalytical Imager.

### 2.12. In vivo imaging

Subcutaneous HepG2 xenografts were established in mice (BALB/c-nu). Chi-GAIN formulations (20 µM in PBS, 60 µL per mouse, tail vein) were administered when tumors reached approximately 100–150 mm^3^. Whole-body near-infrared fluorescence images were acquired under isoflurane anesthesia at the indicated post-injection time points using an *in vivo* imaging system.

### 2.13. Pharmacokinetics

HepG2 tumor-bearing mice received an intravenous tail-vein injection of Chi-GAIN at 5 mg kg^−1^ in sterile PBS. At the indicated post-injection time points, 20 µL of blood was collected from the tail vein into heparinized capillary tubes, immediately centrifuged (3,000*g*, 10 min, 4 °C) to isolate the plasma, and the NIR fluorescence of each plasma sample was measured on an IVIS *in vivo* imaging system. Fluorescence intensities were converted to plasma Chi-GAIN concentrations using a standard curve generated by spiking naïve mouse plasma processed in the same way with known concentrations of Chi-GAIN. Concentration–time data were fitted to estimate the terminal elimination half-life *t*_1/2_.

### 2.14. In vivo PDT

HepG2 tumor-bearing mice were randomized into six groups (*n* = 5 per group): (i) PBS (dark), (ii) PBS + light, (iii) *L*-Chi-GAIN (dark), (iv) *L*-Chi-GAIN + light, (v) *D*-Chi-GAIN (dark), and (vi) *D*-Chi-GAIN **+** light. The mice in the treatment groups received a tail-vein injection of the indicated formulation (20 µM in PBS, 60 µL per mouse). After 24 h, the tumors in the designated “+ light” groups (ii, iv, vi) were exposed to a 730 nm LED at 100 mW cm^−2^ for 20 min (tumor region only; animals under isoflurane anesthesia). The “dark” groups were handled in the same way and shielded from the light. The tumor dimensions (length and width) and body weight were recorded every other day for 14 d after irradiation. The tumor volume *V* was calculated as *V* = (width)^2^ × (length) / 2. After 14 d, the mice were euthanized and the tumors were excised, photographed, and weighed. The TGI was calculated relative to the PBS (dark) control group.

### 2.15. Intratumoral penetration imaging

HepG2 tumor-bearing mice received a single tail-vein injection of *L*-Chi-GAIN or *D*-Chi-GAIN. After 24 h, the mice were euthanized and the tumors excised, embedded in OCT, snap-frozen, and cryosectioned with a thickness of 10 µm. The cryosections were fixed in 4% (w/v) paraformaldehyde in PBS for 20 min at room temperature (∼25 °C), rinsed three times in PBS, and blocked with 10% (w/v) bovine serum albumin for 1 h at 37 °C. Sections were incubated overnight at 4 °C with rat anti-CD31 primary antibody, washed three times in PBS, and incubated for 1 h at room temperature (∼25 °C) in the dark with fluorescein isothiocyanate-conjugated goat anti-rat IgG secondary antibody. The nuclei were counterstained with DAPI, and the sections were washed, mounted with antifade medium, coverslipped, and imaged using fluorescence microscopy.

## 3. Results and Discussion

### 3.1. Galactose induced chiral graphene quantum dots

GQDs were synthesized following our previously established protocol.^36^ Chiral GQDs (*L*- and *D*-GQDs) were then obtained by surface modification with amino-functionalized *L*- and *D*-galactose via *N*-(3-dimethylaminopropyl)-*N*′-ethylcarbodiimide hydrochloride (EDC·HCl)/*N*-hydroxysuccinimide (NHS) coupling (**Fig. 2a**). The zeta-potentials *ζ* of the *L*- and *D*-GQDs shifted to –3.3 ± 2.5 and –3.6 ± 3.0 mV, respectively, indicating that amide formation occurred at the edge carboxyl groups (–COOH) and that the ionization was preserved (**Fig. 2b**). Transmission electron microscopy (TEM) images showed that the *L*- and *D*-GQDs maintained a size distribution of 10–13 nm, which was the same as that of the unmodified GQDs (**Fig. 2c**). High-resolution TEM revealed lattice fringes with a spacing of 0.24 nm, which were attributed to the hexagonal *d*_*1120*_ planes. Unlike the pristine GQDs, multiple interruptions were observed in the lattice fringes of the chiral GQDs, which indicates the presence of lattice distortions and increased out-of-plane curvature. Moreover, the diffuse rings in the electron diffraction patterns indicate that glycosylation induced partial amorphization and increased the severity of the distortion-driven disorder in the GQDs. Atomic force microscopy (AFM) confirmed that the pristine GQDs were less than 0.4 nm thick, which is consistent with monolayer graphene,^43^ whereas the *L*- and *D*-GQDs were more than 0.6 nm thick, reflecting the edge functionalization and increased out-of-plane deformation (**Fig. 2d**). Ultraviolet–visible (UV-vis) spectra showed that *L*- and *D*-GQDs exhibit a red shift of the π→π* band to 258 nm, reflecting the reduction in quantum confinement owing to the electronic coupling between the GQD aromatic lattice and galactose moieties (**Fig. 2e**). In the circular dichroism (CD) spectra, *L*- and *D*-GQDs exhibited negative and positive Cotton effects at 255 and 260 nm, respectively, coincided with their UV-vis absorption signals (**Fig. 2f**). This mirror-image confirms that glycosylation introduced nanoscale distortions into the framework of the GQDs, enabling them with chiroptical activity.^37^ DFT geometry optimizations (B3LYP/6-31G(d), PCM water) of *L*- and *D*-GQD models were further performed to probe nanoscale chirality. Compared with the pristine GQDs, the *L*- and *D*-GQDs exhibited pronounced twisted structures that imparted nanoscale chirality (**Fig. S1**). Reduced density gradient (RDG) analysis revealed abundant intermolecular hydrogen bonds between the galactose and oxygen-containing edge groups of the GQDs, which served as the primary driving force for the observed structural distortion (**Fig. 2g**). Time-dependent DFT (TD-DFT) single-point excited-state calculations using these optimized geometries showed that the Cotton effects in the simulated CD closely matched the experimental spectra (**Fig. S2**). These findings confirmed that chiral GQDs were successfully synthesized via galactosylation, enabling chiral nanostructures.

**Fig. 2.**
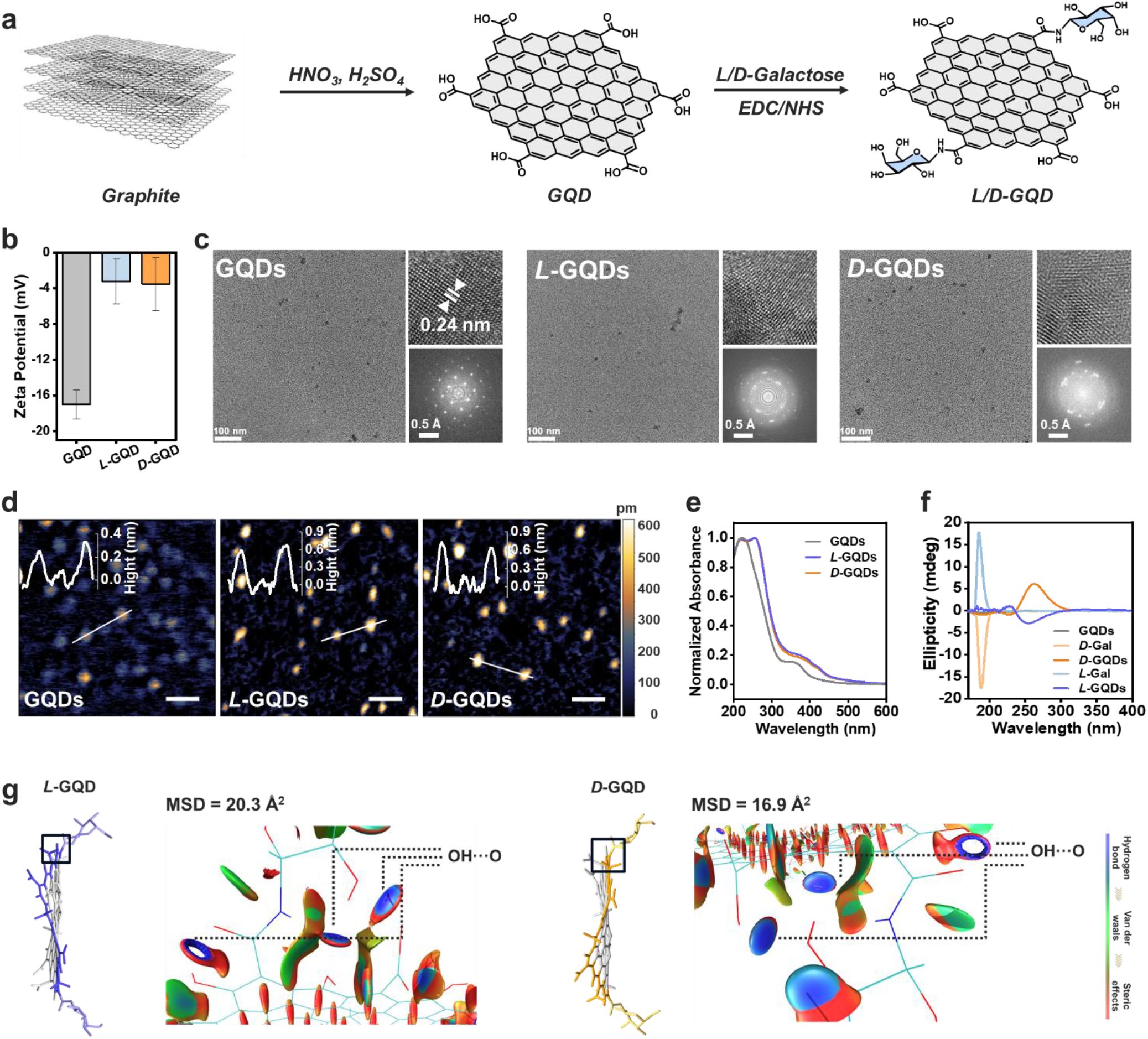
Synthesis and multiscale characterization of chiral GQDs. (a) Molecular schematics showing the synthesis of chiral GQDs. (b) Zeta potentials of GQDs, *L*-GQDs, and *D*-GQDs. (c) Transmission electron microscopy (TEM), lattice alignment, and corresponding fast Fourier-transform images of GQDs, *L*-GQDs, and *D*-GQDs. (d) Atomic force microscopy (AFM) images of GQDs, *L*-GQDs, and *D*-GQDs. Insert: Height profiles along lines. Scale bar: 100 nm. (e) Normalized absorption spectra of GQDs, *L*-GQDs, and *D*-GQDs (20 μM) dispersed in PBS. (f) Circular dichroism (CD) spectra of GQDs, *L*-GQDs, and *D*-GQDs (20 μM). (g) Reduced density gradient (RDG) isosurfaces of the GQDs, *L*-GQDs, and *D*-GQDs with RDG = 0.5, colored based on the sign of *λ*_2_.

### 3.2. Chirality-directed assembly and photophysical modulation for nanocarbon–dye complexes

GQDs, *L*-GQDs, and *D*-GQDs (5 μM) were added to ICG aqueous solution (20 μM) to prepare three nanoassemblies (GAIN, *L*-Chi-GAIN, and *D*-Chi-GAIN, respectively), and their interactions and assembly behaviors were investigated systematically.UV-vis titration experiments showed that *L*-GQD and *D*-GQD significantly reduced the absorption intensity of ICG at 795 nm and induced a new absorption band at 772 nm (**Fig. 3a and 3b**). By contrast, the pristine GQDs only reduced the 795 nm absorption peak and did not generate any new spectral features (**Fig. 3c**). This spectral shift suggests strong exciton coupling between the chiral GQDs and ICG, accompanied by changes in the excited-state properties of ICG.^23^ Fluorescence titration further showed that *L*-GQDs and *D*-GQDs efficiently quenched ICG fluorescence, with Stern-Volmer constants (*K*_SV_) of 4.00 × 10^5^ M^−1^ and 3.80 × 10^5^ M^−1^, respectively, both exceeding that of pristine GQDs and indicating a pronounced nonradiative deactivation pathway. (**Fig. S3**).^44^ TEM images suggested that *L-* and *D-*Chi-GAIN preserved an average diameter of 10-15 nm, with virtually no variation in the particle size distribution or morphology after aging for 14 d (**Fig. S4**). After five sequential 1:2 dilutions in 10% fetal bovine serum (FBS), the absorption profiles remained spectrally invariant, except for the expected attenuation in peak intensity (**Fig. S5**). Chi-GAIN also demonstrated good long-term stability in blood serum, and only a slight deactivation in absorption was observed after storage for 14 d (**Fig. S6**). The assembly behavior was investigated based on an elemental mapping of ICG conducted using energy-dispersive X-ray spectroscopy (EDS) and scanning transmission electron microscopy (STEM) (**Fig. 3e**). ICG was uniformly distributed across the GQD surface in GAIN, whereas it accumulated primarily at the edges in Chi-GAIN. This distinct spatial distribution suggests that the chirality of GQD may affect the site-specific assembly of ICG. urthermore, introducing ICG (2 μM) into concentrated dispersions of *L*-GQD and *D*-GQD (100 μM) led to the formation of larger assemblies, increasing their average particle diameters to approximately 40 nm. By contrast, GQDs without chiral structures formed smaller complexes with a mean size of approximately 18 nm **(Fig. S7)**. This disparity can be attributed to the edge-bound ICG molecules, which promote more pronounced slip stacking between the chiral GQDs via intermolecular interactions. Three distinct ICG/*L*-GQD dimer stacking models (cofacial, parallel-displaced, and head-to-tail stacking) were constructed and geometrically optimized to verify this hypothesis. When ICG was present, hydrogen bond-mediated head-to-tail stacking was thermodynamically preferred over cofacial and parallel-displaced arrangements, whereas the isolated *L*-GQD dimer exhibited the opposite preference (**Fig. S8**). This contrast further corroborated the localized self-assembly of ICG and chiral GQDs. The surface electrostatic potential (ESP) characteristics revealed that the pristine GQD exhibited a high molecular polarity index (MPI) of +10.87 kcal mol^−1^. However, ESP values of the *L*-GQD and *D*-GQD were 5.61 and 5.82 kcal mol^−1^, respectively, suggesting a shift toward edge-localized interaction sites (**Fig. 3f and 3g**). Isothermal titration calorimetry (ITC) confirmed that binding of ICG to chiral GQDs was thermodynamically favorable (Δ*H* ≪0). Notably, the association constant *K*_a_ for the chiral GQDs was significantly higher than that for the pristine GQDs (**Fig. 3h–3j and S9**). These results indicate that chiral GQDs modulate the assembly pattern and photophysical properties of ICG by enhancing the site-specific electrostatic interactions at the edges. Cyclic voltammetry was used to determine the reduction potential *E*_red_ of GQDs and the oxidation potential *E*_ox_ of ICG to assess the possibility of photoinduced intermolecular electron transfer (**Fig. 3k and S10**). The change in the Gibbs free energy Δ*G* was estimated using the Rehm–Weller equation, which confirmed that electron transfer from ICG to chiral GQDs is thermodynamically favorable (**Fig. 3l and S11**). Together, these findings indicate that chiral GQDs drive the directed self-assembly of ICG, producing stable Chi-GAIN nanocomplexes with efficient electron transfer capabilities (**Fig. 3m**).

**Fig. 3.**
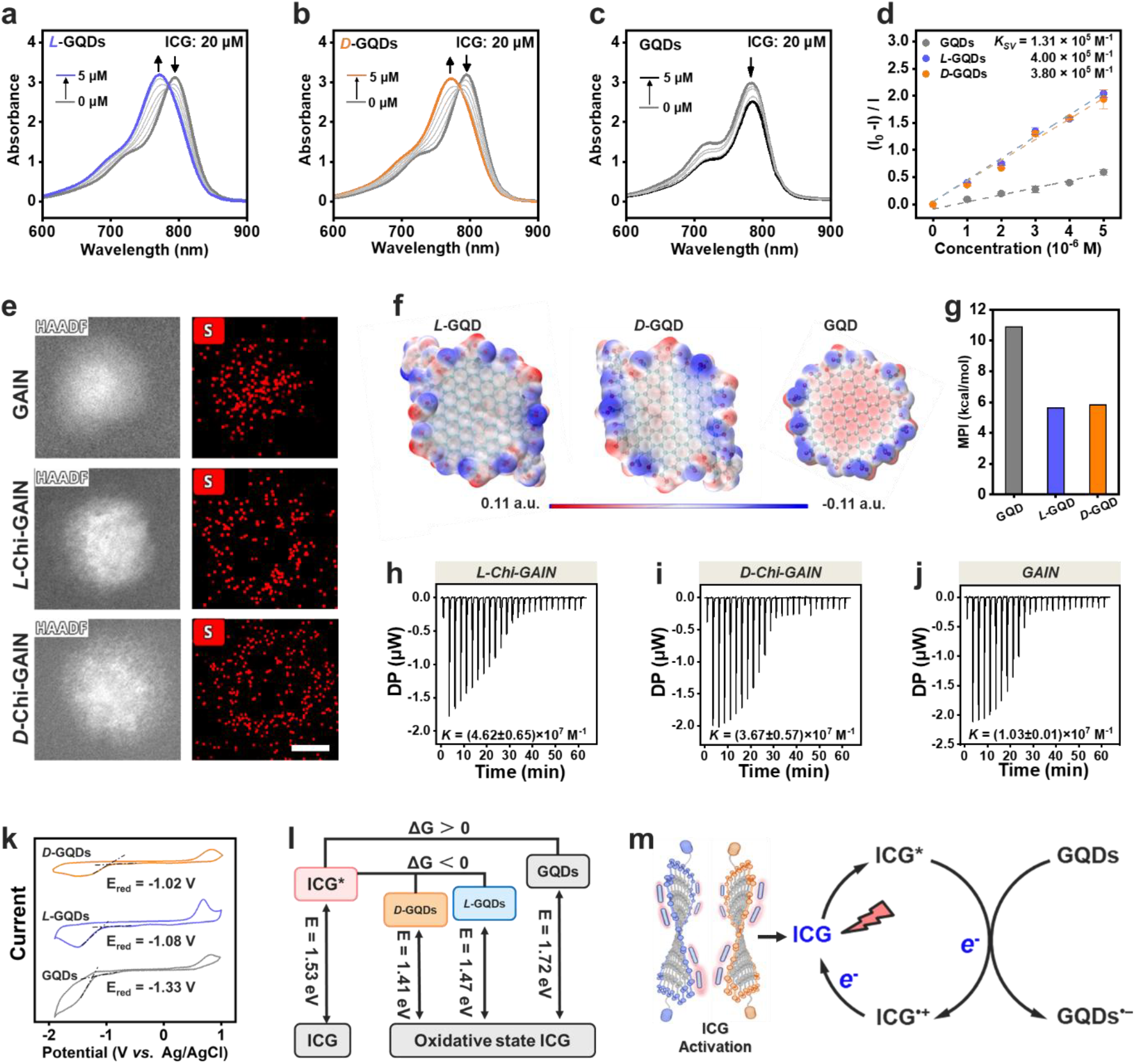
Mechanism of directional self-assembly of Chi-GAIN with photoinduced electron transfer ability. (a-c) Absorption spectra of ICG in *L*-GQD, *D*-GQD, and GQD solutions with different concentrations. (d) Stern-Volmer plots for fluorescence quenching of ICG by *L*-GQDs, *D*-GQDs, and GQDs in PBS. (e) Scanning transmission electron microscopy–high-angle annular dark field (STEM HAADF) images of GAIN, *L*-Chi-GAIN, and *D*-Chi-GAIN and the corresponding sulfur distributions. Scale bar: 5 nm. (f) Electrostatic potential surfaces of *L*-GQD, *D*-GQD, and GQD. (g) Molecular polarity indices (MPI) of *L*-GQD, *D*-GQD, and GQD. (h-j) Isothermal titration calorimetry (ITC) of *L*-GQDs, *D*-GQDs, and GQDs (10 μM) titrated with ICG (20 μM) at 298 K in PBS. (k) Cyclic voltammetry curves of *L*-GQDs, *D*-GQDs, and GQDs in water containing 0.1 M KCl with Ag/AgCl as the reference electrode, a glassy carbon electrode as the working electrode, and a Pt wire as the counter electrode; scan rate, 100 mV s^−1^. (l) Gibbs free energy of the electron transfer between ICG and different GQDs. (m) Schematic of the photoinduced charge transfer between ICG and chiral GQDs.

### 3.3. Chi-GAIN–driven Type-I ROS generation under hypoxia

Then, dihydroethidium (DHE) and 9,10-anthracenediyl-bis(methylene)dimalonic acid (ABDA) were used to distinguish between Type-I and Type-II ROS. The absorbance of ABDA remained approximately constant under irradiation in the presence of all three nanocomplexes (GAIN, *L*-Chi-GAIN, and *D*-Chi-GAIN), indicating negligible generation of singlet oxygen (^1^O_2_) via the Type-II pathway (**Fig. S12**). The potential of the nanocomplexes to engage in Type-I photoreactions was investigated by using NADH, a central biological reductant and key electron/proton donor in cellular metabolism, as a model substrate. Upon irradiation, the DHE fluorescence in the *L*-Chi-GAIN and *D*-Chi-GAIN solutions containing NADH increased significantly, suggesting that O_2_^•–^ was produced (**Fig. 4a, 4b and S13**). The photocatalytic activity of Chi-GAIN toward NADH oxidation was verified by using ^1^H nuclear magnetic resonance (NMR) spectroscopy to monitor the conversion of NADH to NAD^+^. In the presence of Chi-GAIN under irradiation, new peaks corresponding to NAD^+^ appeared, providing direct evidence of the photooxidation reaction (**Fig. 4c**). By contrast, no NAD^+^ signals were observed with GAIN under the same conditions (**Fig. S14)**. The photocatalytic efficiency of Chi-GAIN for NADH oxidation in aqueous solution was further assessed using UV-vis spectroscopy. In the presence of irradiation, the characteristic absorption of NADH at 339 nm decreased significantly, whereas it remained unchanged without irradiation **(Fig. S15)**. This confirms that NADH is effectively photooxidized by Chi-GAIN. Notably, *L*-Chi-GAIN and *D*-Chi-GAIN achieved turnover frequencies of 49.8 and 47.0 min^−1^, respectively, which were between two and three orders of magnitude higher than those reported for commonly studied transition-metal catalysts **(Fig. S16)**.^45, 46^ Electron spin resonance (ESR) spectroscopy with 5,5-dimethyl-1-pyrroline-N-oxide (DMPO) spin trapping revealed the characteristic DMPO–O_2_^•–^ quartet from photo-excited Chi-GAIN (**Fig. 4d**), whereas the irradiated GAIN produced no detectable radical signal. This confirms that GQD chirality is indispensable for modulating the excited-state energetics of ICG and enabling efficient O_2_^•–^ generation. This mechanism was investigated by computing the frontier molecular orbitals at the B3LYP/6-31G(d) level (**Fig. 4e**). For free ICG, the highest and lowest occupied molecular orbitals (HOMO and LUMO, respectively) were localized on the polymethine bridge. Within the Chi-GAIN assembly, ICG and chiral GQD formed a stable charge-transfer complex that spatially separated the HOMO from the LUMO, thereby facilitating more efficient photoinduced charge separation. The HOMO-LUMO gaps *E*_*g*_ of ICG, *L*-Chi-GAIN, and *D*-Chi-GAIN were 2.07, 2.25, and 2.22 eV, respectively, which is consistent with the slight blue shift in the absorption spectrum. The excited-state energy gaps of ICG, *L*-Chi-GAIN, and *D*-Chi-GAIN were also evaluated (**Fig. 4f**). For free ICG, ISC to T_1_ was inefficient because the *S*_*1*_*–T*_*1*_ gap was relatively large (0.28 eV). By contrast, this gap was only 0.18 eV for *L*-Chi-GAIN and 0.17 eV for *D*-Chi-GAIN. Therefore, chiral GQDs increase the ISC of ICG through excitonic coupling and serve as effective electron acceptors, which promotes excited-state electron transfer and increases the O_2_^•–^ generation efficiency (**Fig. 4g**).

**Fig. 4.**
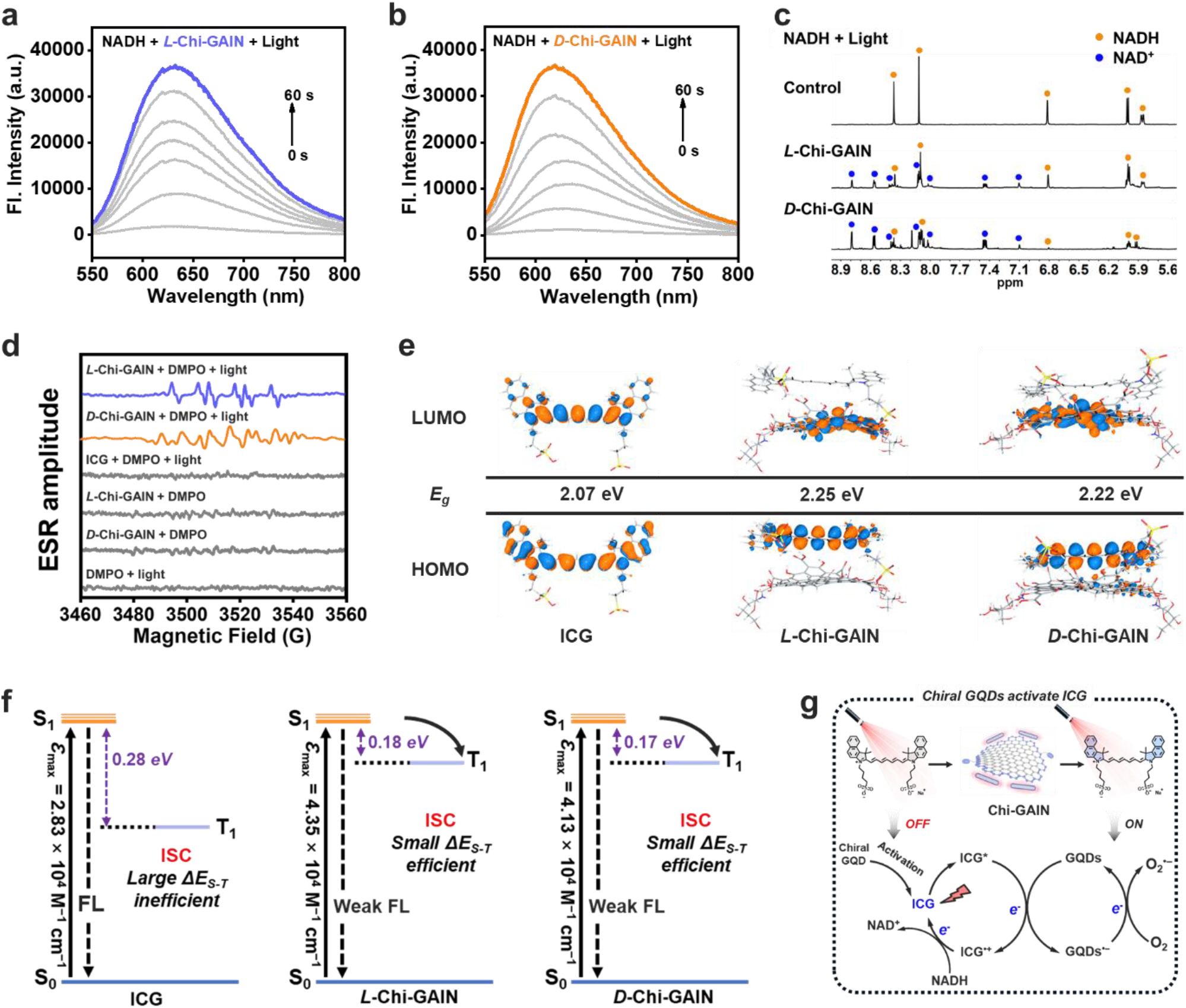
Excitonic coupling boosts ISC and Type-I ROS generation in Chi-GAIN. (a,b) Fluorescence spectra of dihydroethidium (DHE) (50 μM) containing 500 μg mL^−1^ of ctDNA after irradiation (730 nm, 20 mW cm^−2^) for different durations in the presence of *L*-Chi-GAIN and *D*-Chi-GAIN (10 μM) dispersed in phosphate-buffered saline (PBS). (c) ^1^H nuclear magnetic resonance (NMR) of β-nicotinamide adenine dinucleotide (reduced form, NADH) in D_2_O after different treatments. (d) Electron spin resonance (ESR) spectra to detect the O_2_^•–^generated by Chi-GAIN under irradiation using 5,5-dimethyl-1-pyrroline-N-oxide (DMPO) as a spin-trap agent. (e) Frontier molecular orbitals determined from time-dependent density functional theory (TD-DFT) calculations of ICG, *L*-Chi-GAIN, and *D*-Chi-GAIN in their ground states (S_0_). (f) Jablonski diagrams of ICG, *L*-Chi-GAIN, and *D*-Chi-GAIN. (g) Schematic showing photoinduced electron transfer and ROS generation

### 3.4. PDT–triggered lysosomal rupture, pyroptosis, and immunogenic cell death pathways

Galactose-functionalized GQDs can selectively bind to ASGPR, which is overexpressed on HCCs, enabling receptor-mediated endocytosis through clathrin-coated pits.^47^ This dual role enhances both cellular uptake and lysosomal trafficking, and simultaneously promotes chirality-dependent biointeractions with biological systems.^48^ Confocal laser scanning microscopy (CLSM) results showed that *L-* and *D-*glycosylation both promoted GQD internalization in human hepatocellular carcinoma (HepG2) cells, primarily through selective binding to ASGPR. Notably, the overall uptake of *D*-Chi-GAIN exceeded that of *L*-Chi-GAIN (**Fig. S17**). Moreover, even when ASGPR was blocked with a specific antibody, the HepG2 cells still internalized more *D*-Chi-GAIN than *L*-Chi-GAIN (**Fig. S18**). The same preference was also observed at 4 °C, a temperature that largely suppresses energy-dependent endocytic pathways, indicating that *D*-Chi-GAIN traverses the lipid membrane more efficiently via passive diffusion. This is consistent with previous studies, which have shown that *D*-GQDs exhibit enhanced transmembrane transport through passive diffusion compared to their *L*-counterparts.^36, 37^ Pearson correlation coefficients for *L*-Chi-GAIN and *D*-Chi-GAIN with LysoView™594 were 0.84 and 0.87, respectively, indicating effective lysosomal targeting by glycosylated GQDs (**Fig. 5a**). Next, we used the general ROS probe 2′,7′-dichlorodihydrofluorescein diacetate (DCFH-DA) to assess the ability of Chi-GAIN to generate ROS under normoxicand hypoxic conditions. Under both normoxic and hypoxic conditions, strong intracellular green fluorescence was observed upon irradiation. Therefore, Chi-GAIN efficiently generated ROS and was essentially insensitive to oxygen tension (**Fig. 5b**). Consistent with this observation, HepG2 cells treated with Chi-GAIN and DHE displayed a characteristic red fluorescence after light exposure, confirming the formation of O_2_^•–^. Furthermore, the fluorescence of hydroxyphenyl fluorescein (HPF), a hydroxyl-radical (^•^OH) sensor, increased substantially under the same conditions, indicating that the photogenerated O_2_^•–^ underwent a cascade reaction to yield ^•^OH, which can inflict oxidative damage on cells. Quantitative NADH depletion assays using WST-8 demonstrated a significant reduction in NADH proportional to the concentrations of *L*-Chi-GAIN and *D*-Chi-GAIN upon irradiation, and negligible reduction without irradiation (**Fig. 5c**). Cytotoxicity assays using Cell Counting Kit (CCK)-8 revealed pronounced cytotoxicity under irradiation for both *L*-Chi-GAIN and *D*-Chi-GAIN, with IC_50_ values of 12.32 and 7.23 µg mL^−1^, respectively, which were consistent across normoxic and hypoxic conditions (**Fig. 5d**). By contrast, the cellular phototoxicity of the commercial Type-II PS chlorin e6 (Ce6) was significantly suppressed under hypoxic conditions (**Fig. S19**). These findings indicate that Chi-GAIN, which functions as a Type-I PS, shows substantially lower dependence on oxygen availability than conventional PSs, and can inhibit tumor cell proliferation even in hypoxia. Under both conditions, minimal dark toxicity was observed, as corroborated by Calcein-AM and propidium iodide (PI) staining (**Fig. S20**).

**Fig. 5.**
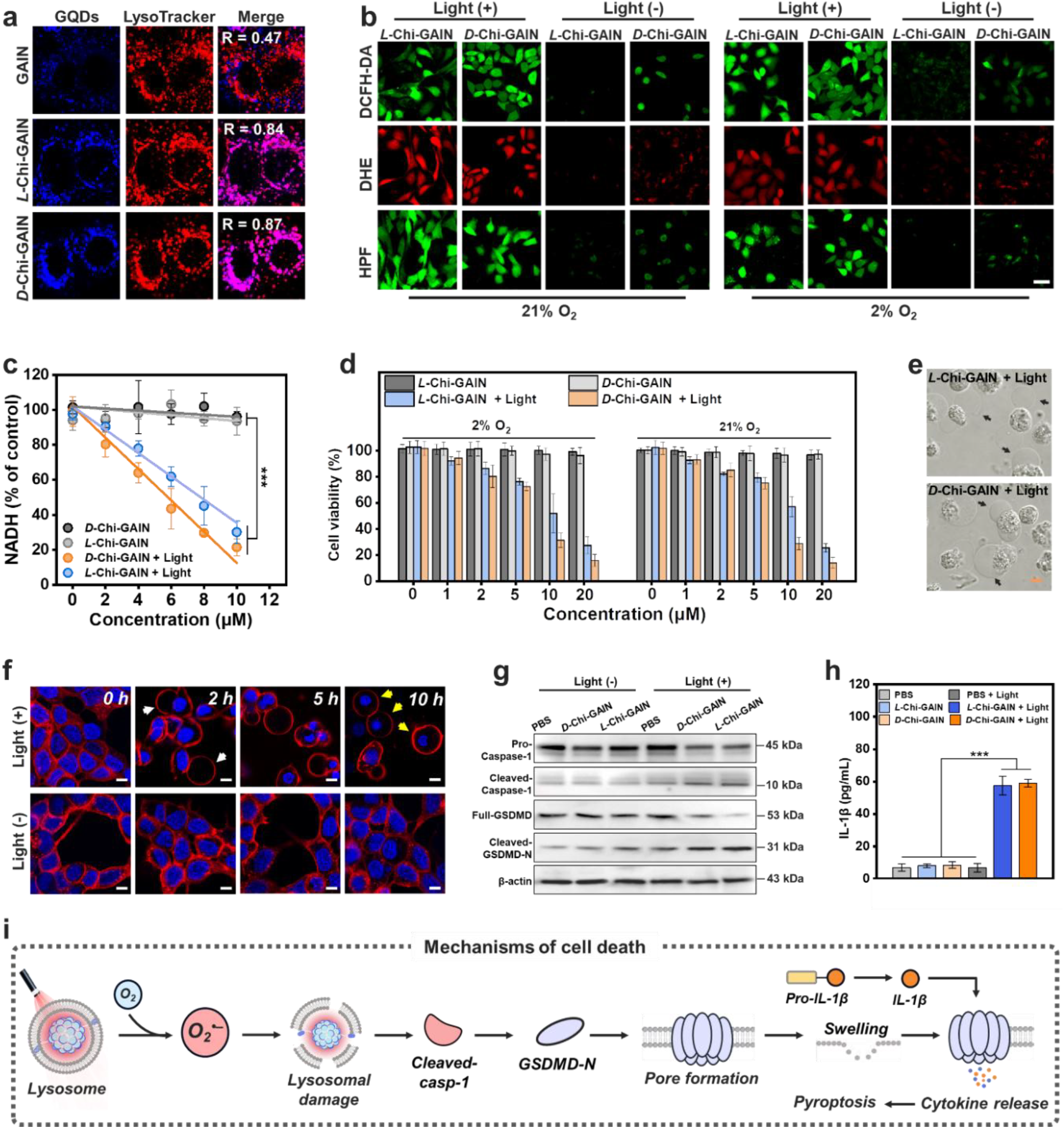
Chi-GAIN inducing lysosomal disruption and Caspase-1-mediated pyroptosis. (a) Co-localization of GAIN, *L*-Chi-GAIN, and *D*-Chi-GAIN with LysoTracker in human hepatocellular carcinoma (HepG2) cells after incubation for 4 h. Scale bar: 10 μm. (b) ROS imaging under normoxia (21% O_2_) or hypoxia (2% O_2_) using DCFH-DA (general ROS), DHE (O_2_^•–^), and hydroxyphenyl fluorescein (HPF) (^•^OH) probes. Scale bar: 50 µm. (c) Intracellular NADH levels after treatment with increasing Chi-GAIN concentrations (*n* = 3). (d) Viability of HepG2 cells subjected to various concentrations of Chi-GAIN in the absence or presence of light-irradiation (730 nm, 10 mW cm^−2^, 10 min) under normoxic or hypoxic conditions (*n* = 5). (e) Morphological features of HepG2 cells treated with Chi-GAIN in the presence of irradiation. Black arrows represent pyroptotic bodies. Scale bar: 10 μm. (f) Bubble-like protrusions in the HepG2 cell membrane and nuclear shrinkage at different time points after irradiation. 1,1′-dioctadecyl-3,3,3′,3′-tetramethylindodicarbocyanine (DiD) was used as a membrane marker, and Hoechst 33342 was used as a nuclear marker. Scale bar: 10 μm. (g) Cleavage of Gasdermin-D (GSDMD) and caspase-1 from HepG2 cells under different treatment conditions, as detected by western blot analysis. (h) Quantification of IL-1β released from HepG2 cells after different treatments (*n* = 5). (i) Schematic showing the mechanisms of O_2_^•–^-induced lysosomal disruption and caspase-1-mediated pyroptosis. Data are expressed as the mean ± standard deviation (SD). Statistical differences were analyzed using the Student’s two-sided *t*-test. \*\*\**P* < 0.001.

We investigated whether Chi-GAIN could disrupt lysosomes and induce Gasdermin-D (GSDMD)-mediated pyroptosis under irradiation by monitoring the lysosomal integrity using acridine orange (AO). HepG2 cells exposed to Chi-GAIN in the absence of irradiation exhibited intense, punctate red fluorescence, indicating that the lysosomes remained intact (**Fig. S21)**. Irradiation virtually extinguished this red signal in the Chi-GAIN group, confirming that photoactivated Chi-GAIN elicits pronounced lysosomal disruption. Notably, the cells treated with Chi-GAIN and then irradiated developed conspicuous bubble-like protrusions (**Fig. 5e**). Analysis of cell morphology showed that these membrane blebs enlarged progressively, whereas the nucleus gradually shrank, although it remained intact. These signs closely match those of pyroptotic cell death (**Fig. 5f**).^49^ Western blot analysis revealed that, under irradiation, HepG2 cells treated with *L*-Chi-GAIN and *D*-Chi-GAIN showed markedly increased cleavage of GSDMD-N and caspase-1 (**Fig. 5g**). Moreover, membrane pores formed by GSDMD facilitated the release of lactate dehydrogenase and inflammatory cytokines (IL-1*β* and IL-18) (**Fig. 5h and S22**). These findings show that Chi-GAIN can function as an efficient O_2_^•–^generator, triggering lysosomal disruption and caspase-1-mediated pyroptosis (**Fig. 5i**).

### 3.5. Chirality-driven HA modulation for enhanced intratumoral penetration

Given the excellent performance of Chi-GAIN in cellular studies, we evaluated its tumor penetration efficiency *in vivo*. First, a HepG2 tumor-bearing BALB/c mouse model was established via subcutaneous injection. Upon intravenous administration, Chi-GAIN rapidly accumulated at the tumor site, reaching peak fluorescence intensity after 24 h. The fluorescence signal in the major organs was eliminated within 24 h, indicating minimal off-target retention (**Fig. 6a and S23**). Notably, *L*-Chi-GAIN exhibited prolonged retention at the tumor site for up to 48 h, which was significantly longer than that for *D*-Chi-GAIN. Moreover, pharmacokinetic analysis showed that *L*-Chi-GAIN and *D*-Chi-GAIN extended the plasma half-life *t*_*1/2*_ of ICG by factors of 4.9 and 3.8, respectively, expected to improve tumor accumulation and overall therapeutic efficacy (**Fig. S24**). We investigated whether Chi-GAIN enhanced deep intratumoral penetration by performing immunohistochemistry (IHC) staining on isolated tumors (**Fig. 6b**).^50^ At 24 h post-injection, most of the *D*-Chi-GAIN remained within approximately 150 µm of the vasculature, whereas the *L*-Chi-GAIN penetrated beyond 350 µm into the tumor, confirming its superior intratumoral penetration. Quantitative colocalization analysis yielded high Pearson coefficients between the GQD and ICG channels, indicating that the spatial distribution of ICG closely depended on the penetration capability of the chiral GQD (**Fig. S25**). Because most PSs are delivered via biocompatible nanoliposomes, we also compared two commercial formulations (DPPC and DSPE-PEG_2000_) with Chi-GAIN and assessed intratumoral penetration *in vivo*. Fluorescence imaging revealed that ICG delivered by both liposomal carriers remained almost exclusively in perivascular regions, with penetration depths of only ∼120 µm. This finding highlights the challenge posed by the dense interstitial tumor matrix, which acts as a diffusion barrier that has hindered deep drug penetration for nearly two decades.^51, 52^ Collectively, these results demonstrate that constructing chiral nanocarriers via glycosylation is an effective strategy to enhance both tumor targeting and penetration. To elucidate the penetration mechanism of *L*-Chi-GAIN, we performed Molecular dynamics (MD) simulations to compare the interactions of chiral GQD with a collagen-HA ECM model. Under a constant 50 pN longitudinal drag force (convective interstitial flow), *L*-GQD exhibited an average axial velocity approximately 3.7-fold higher than that of *D*-GQD (**Fig. 6c and S26**). This greater displacement was associated with significantly fewer adhesive contacts between *L*-GQD and the surrounding ECM (**Fig. 6d**). Notably, in unbiased simulations from random placements, the fraction of frames with close-range HA pairing was 28.5% for *L*-GQD and 65.7% for *D*-GQD, consistent with shorter residence times and a shallower GQD-HA energy minimum for *L*-GQD (**Fig. 6e and 6f**). Subsequently, ITC corroborated this stereoselectivity: thermograms for HA titrations into *D*-GQD displayed consistently larger injection heats than those for *L*-GQD. Global one-site fits returned higher *K*_a_, more exothermic Δ*H*, and a significantly higher binding stoichiometry (*N*) for *D*-GQD, indicating stronger and more multivalent engagement of HA (**Fig. 6g and 6h**). Given that HA is a primary diffusion barrier in tumor stroma, low HA affinity of *L*-GQD facilitates diffusion through the ECM, thereby limiting non-specific matrix sequestration and providing a mechanistic basis for the deep intratumoral penetration and prolonged tumor retention observed i*n vivo*.

**Fig. 6.**
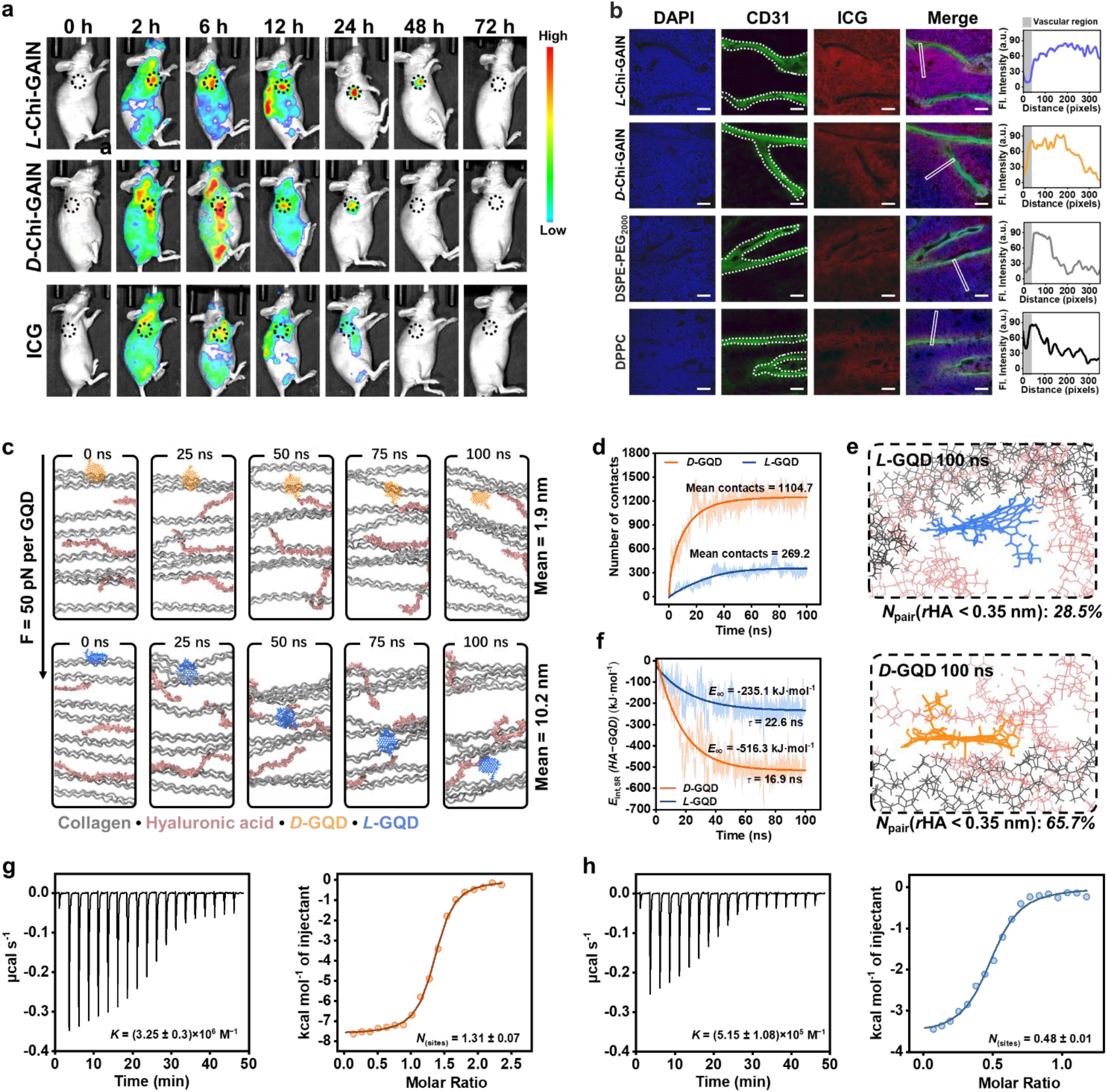
Tumor penetration effect of *L*-Chi-GAIN of and potential mechanisms. (a) *In vivo* fluorescence imaging of Chi-GAIN and ICG at different time points after intravenous injection (i.v.). (b) Fluorescence images showing the penetration of various ICG delivery systems into deep tumor tissues. Blue: nucleus; green: blood vessels; red: ICG. Scale bar: 200 μm. (c) Representative simulation snapshots of *L*- and *D*-GQD penetrating through a model extracellular matrix (ECM) under a constant 50 pN pulling force at different times. (d) The number of atomic contacts between *L*- or *D*-GQDs and the ECM during the force-induced infiltration at different times. (e) Representative snapshots of *L*- and *D*-GQDs in the ECM model under unbiased (no-force) conditions after 100 ns. (f) GQD-HA interaction energy profiles from unbiased MD simulations. (g-h) ITC of *D*-GQD and *L*-GQD (10 μM) titrated with HA (200 μM) at 298 K in PBS.

### 3.6. Enhanced in vivo PDT via Chi-GAIN through hypoxia mitigation and barrier modulation

The *in vivo* PDT performance of Chi-GAIN was assessed using HepG2 tumor-bearing mice, which were divided into six groups: PBS, PBS + light, *L*-Chi-GAIN, *L*-Chi-GAIN + light, *D*-Chi-GAIN, and *D*-Chi-GAIN + light. Guided by *in vivo* fluorescence imaging, irradiation was performed 24 h post-injection (**Fig. 7a**). The body weights of the mice increased slightly during the subsequent 14 d (**Fig. 7b**), and the tumor growth in the group treated with Chi-GAIN + light was significantly inhibited compared to that in the PBS group (**Fig. 7c**). The mice were euthanized after 14 d, and all the tumor tissues were excised and weighed. The results showed that *D*-Chi-GAIN achieved tumor-growth inhibition (TGI) of 68.8%, whereas *L*-Chi-GAIN blocked tumor growth almost completely, achieving TGI of 98.2% (**Fig. 7d and 7e**). The latter TGI is approximately 2.5 times that of the hepatocellular-carcinoma drug sorafenib; therefore, *L*-Chi-GAIN exhibits superior antitumor efficacy *in vivo*.^53^ Unlike many FDA-approved PSs that require long drug-light intervals and extra illumination or repeat dosing, this therapy achieves marked tumor inhibition with a single administration and 20 min irradiation, sharply shortening the treatment window and highlighting clinical potential.^54^ This enhanced performance is attributed to its efficient deep intratumoral penetration and significantly improved PDT efficacy via Type-I ROS generation. Finally, the biosafety and antitumor efficacy of Chi-GAIN were evaluated using hematoxylin and eosin (H&E) staining (**Fig. 7f and S27**). Extensive tumor cell death and tissue damage were observed in the treated groups, whereas no histopathological abnormalities were detected in the major organs. This confirms that Chi-GAIN is safe and effective for *in vivo* PDT.

## 4. Conclusion

In summary, this study developed a chiral GQD-ICG nanoassembly (*L*-Chi-GAIN) that can effectively address the critical challenges inherent in PDT, particularly tumor hypoxia and limited tissue penetration. The introduction of nanoscale chirality through the surface galactosylation of GQDs induces stereogenic structural distortion, thereby reducing the molecular polarity and promoting the site-selective assembly of ICG. This strategic self-assembly significantly increases the photoinduced electron transfer efficiency and O_2_^•–^generation. More importantly, nanoscale chirality modulates GQD-ECM interactions, with *L*-GQDs showing less close-range HA pairing and a shallower GQD-HA energy minimum, which together reduce stromal retention and enhance interstitial transport. Under NIR irradiation, *L*-Chi-GAIN induces robust oxidative stress, resulting in lysosomal disruption and triggering Gasdermin D-mediated pyroptosis in cancer cells. Overall, our findings highlight the significant potential of incorporating nanoscale chirality into nano-PS designs, offering a generalizable design strategy to overcome oxygen limitation and stromal diffusion barriers.

**Fig. 7.**
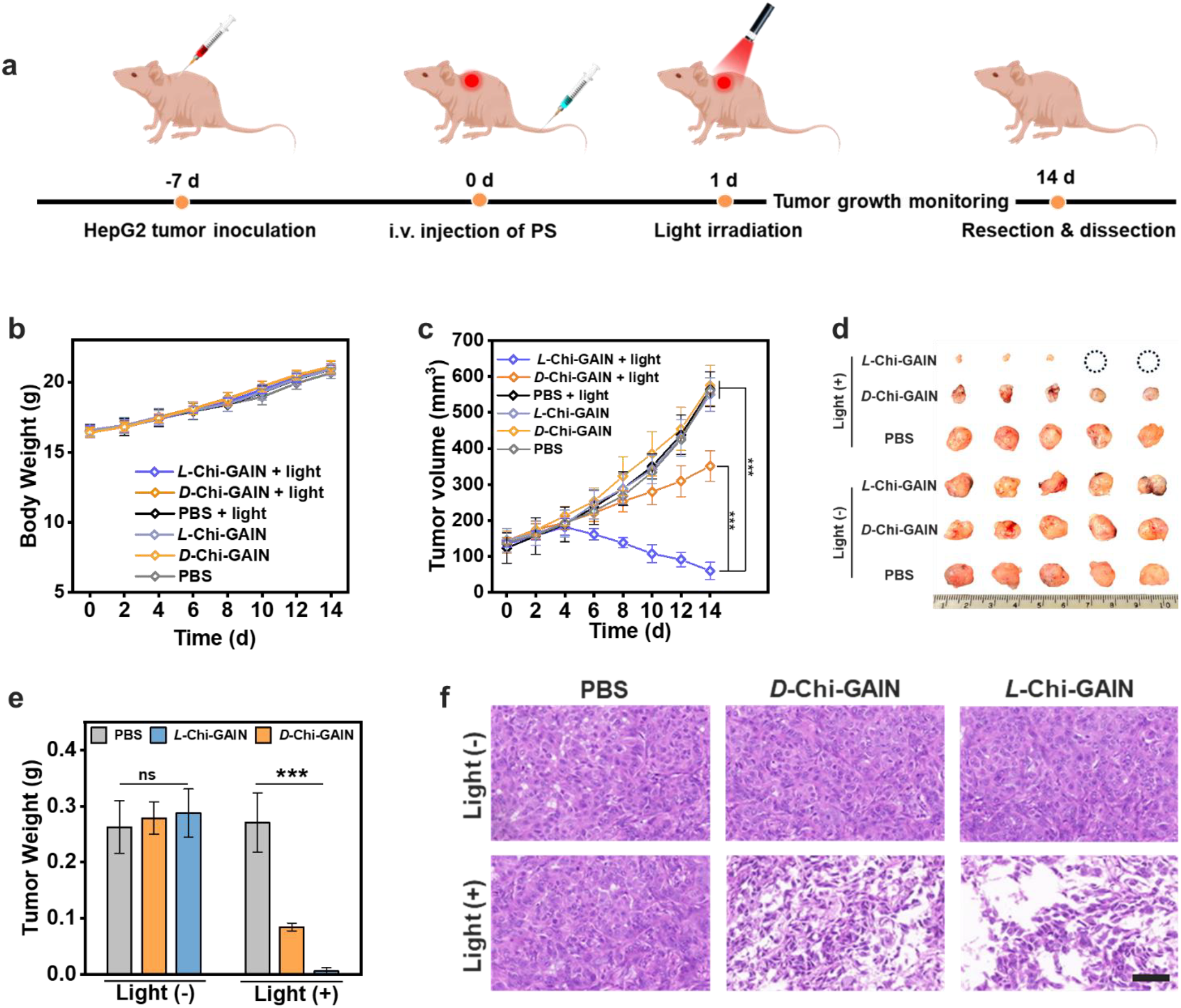
In vivo PDT effects of *L*-Chi-GAIN and *D*-Chi-GAIN. (a) Schematic showing the tumor implantation schedule, Chi-GAIN injection, irradiation, and tumor growth monitoring. (b) Changes in body weight of the mice after different treatments (*n* = 5). (c) Tumor volumes of mice after different treatments (*n* = 5). (d) Images of tumor tissues from different groups of tumor-bearing mice. (e) The weight of the tumors in the different groups after 14 d (*n* = 5). (f) Hematoxylin and eosin (H&E) staining of tumor tissues collected from the mice in each group. Scale bar: 50 μm. Data are expressed as the mean ± SD. Statistical differences were analyzed using the Student’s two-sided *t*-test. \*\*\**P* < 0.001.

## Supporting information

support information

## Authorship contribution

**Yichun Wang**: Conceptualization, Supervision, Funding acquisition, Writing – review & editing. **Yichen Liu**: Conceptualization, Investigation, Formal analysis, Writing – original draft. **Gaeun Kim**: Investigation, Formal analysis. **Runyao Zhu**: Investigation, Formal analysis. **Hyunsu Jeon**: Investigation, Formal analysis.

## Acknowledgements

This work was supported by the NIH MIRA (NIH 1R35GM15608-01) and the NSF Career Award (NSF CBET-2337387). This research was funded in part by the BELS Supplemental Professional Development Award (Bioengineering and Life Sciences Initiative, University of Notre Dame; Grant No. 374553 28016). The TEM and CLSM images were obtained using the instruments at the Notre Dame Integrated Imaging Facility (NDIIF). The Western blot images and ITC experimental data were acquired using the instrument at the Biophysics Instrumentation Core (BIC), University of Notre Dame. Computational work was performed using the high performance computing clusters at the Center for Research Computing (CRC), University of Notre Dame. The mice were bred and maintained under specific pathogen-free conditions at the Freimann Life Science Center (FLSC), University of Notre Dame. We appreciate the support provided by these facilities.

## Data Availability

Data will be made available on request.

