## Supplementary material for "Nanochirality-programmed Type-I photosensitizer enables deep-tumor photodynamic therapy by reducing extracellular-matrix adhesion": support information

**Materials and Instruments**

Materials and reagents were purchased from commercial suppliers and used without further purification. All solvents were analytical pure unless otherwise noted. Distilled water was used throughout the experiments. Absorption spectra of liquid samples were determined on JASCO V-670 UV-Vis/NIR Spectrophotometer. The fluorescence spectra were recorded by an Infinite M1000 plate reader (Tecan). CD spectra were recorded on a CD spectroscopy (Jasco J-1700 Spectrometer). Cyclic voltammetry (CV) experiments were carried out at room temperature with a potentiostat (Autolab PGSTAT302 N, Metrohm Autolab B.V., Utrecht, The Netherlands). Transmission electron microscope (TEM) images were obtained using a FEI Talos F200S instrument. NMR spectra were recorded with Bruker AVANCE III HD 400 Nanobay. The zeta potentials were recorded using Malvern Zetasizer Nano ZS (Malvern Panalytical; Worcestershire, United Kingdom). Confocal fluorescence imaging was performed with Nikon A1R MP multiphoton microscopy. Irradiation was performed using a High power LED lamp source (CL-1501, Asahi Spectra Co., Ltd., Tokyo, Japan). Atomic Force Microscope (AFM) images were recorded using Jupiter XR AFM. In vivo imaging was recorded using an AMI HT optical imaging system (Advanced Molecular Imager High Throughput, Spectral Instruments Imaging, Tucson, AZ, USA). Isothermal titration calorimetry (ITC) experiments were carried out on a MicroCal PEAQ-ITC calorimeter (Malvern Panalytical Ltd., Malvern, UK).

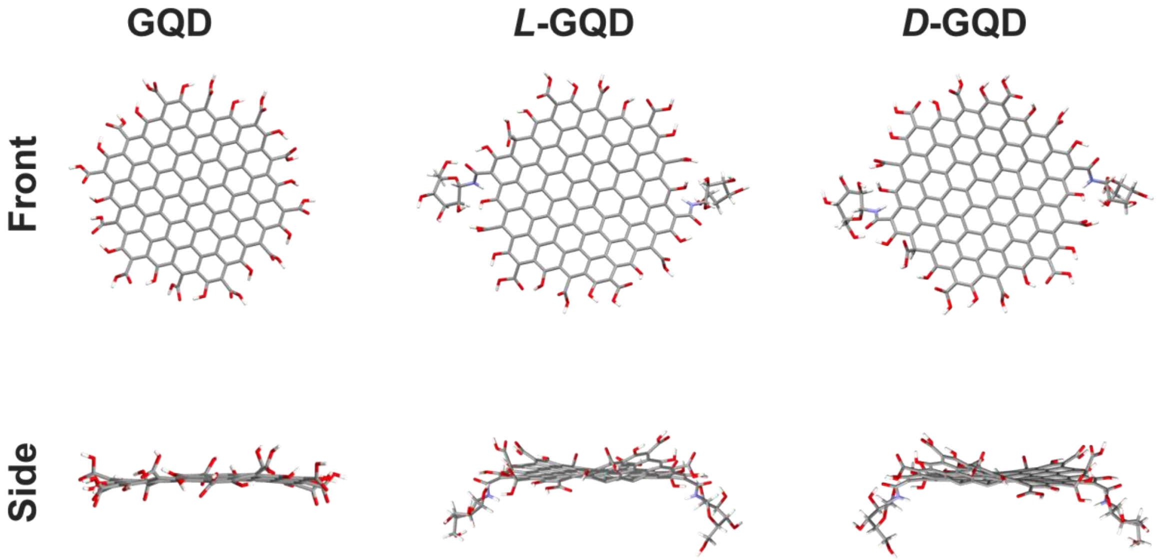


**Figure S1.** Optimized geometries of GQDs, L-GQDs, and D-GQDs at the B3LYP/6-31G(d) level..

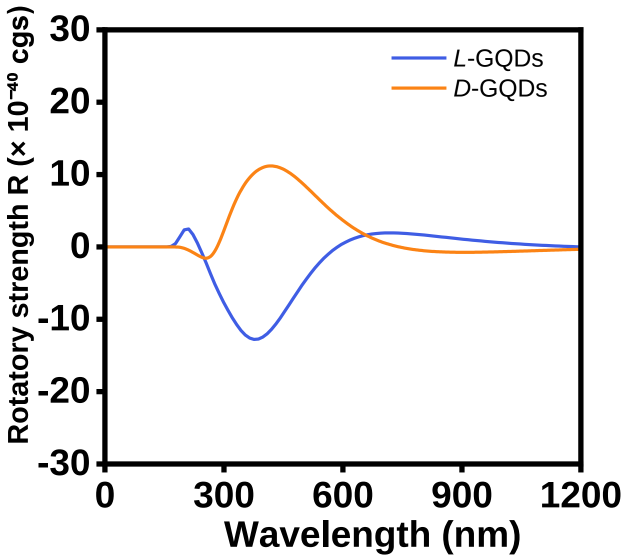


**Figure S2.** Simulated electronic circular dichroism (ECD) spectra (rotatory strength, R × $10^{-40}$ cgs) of chiral graphene quantum dots (GQDs) (*L*-GQD and *D*-GQD) calculated at the TD-B3LYP/6-31G(d) level; spectral lines obtained by Gaussian broadening with 0.25 eV HWHM.

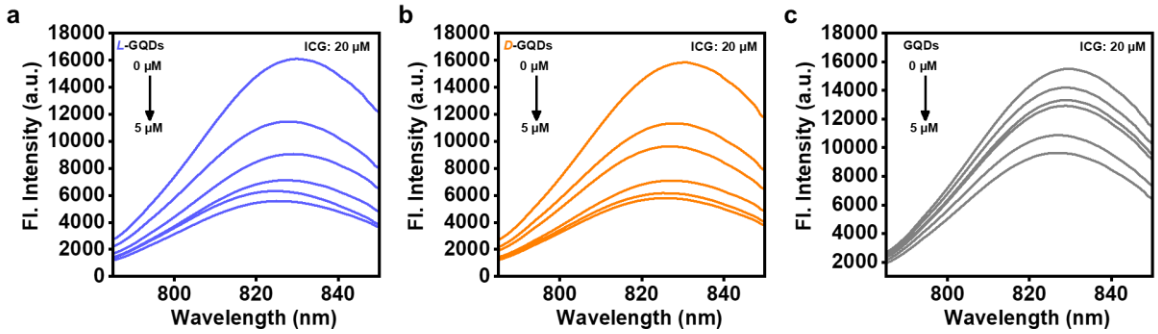


**Figure S3.** The fluorescence emission spectra of Indocyanine Green (ICG) (2.0 μM) phosphate-buffered saline (PBS) with increasing amounts of (a) ***L*-GQDs**, (b) ***D*-GQDs** and (c) **GQDs** (0–5 μM).

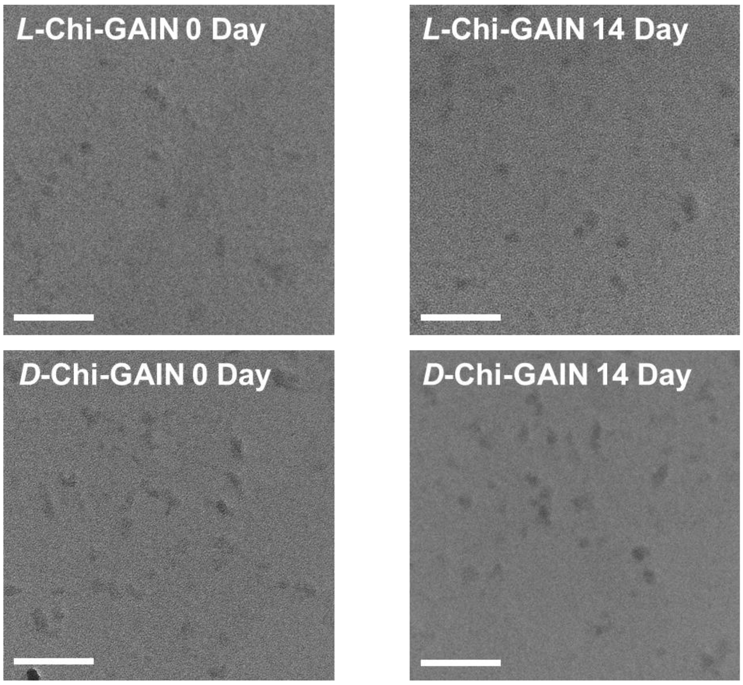


**Figure S4.** Transmission electron microscopy (TEM) image of ***L*-Chi-GAIN** and ***D*-Chi-GAIN** (**Chi**ral **G**r**a**phene quantum dot–**I**ndocyanine Green **N**anoassembly) in water after 0 day and 14 days storage. Scale bar: 50 nm.

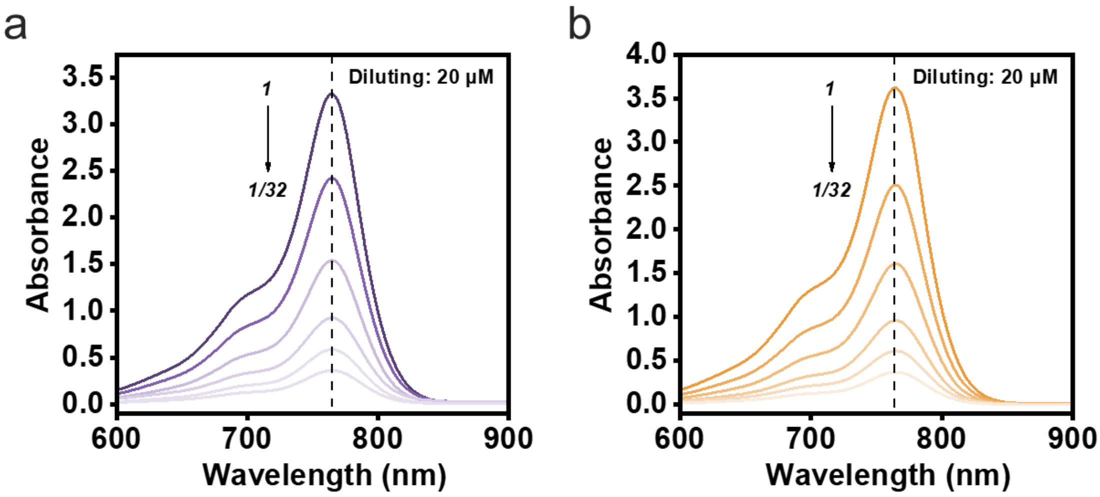


**Figure S5.** Absorption spectra of (a) ***L*-Chi-GAIN** and (b) ***D*-Chi-GAIN** (20 μM) in PBS with 10% Fetal bovine serum (FBS), and after gradual dilutions.

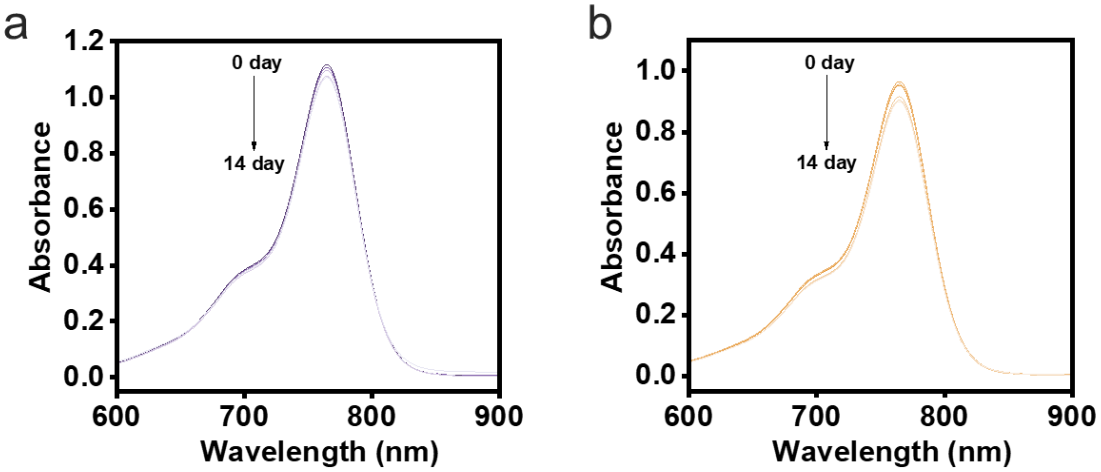


**Figure S6.** Absorption spectra of (a) ***L*-Chi-GAIN** and (b) ***D*-Chi-GAIN** (2 μm) in PBS with 10% FBS after being stored for different times.

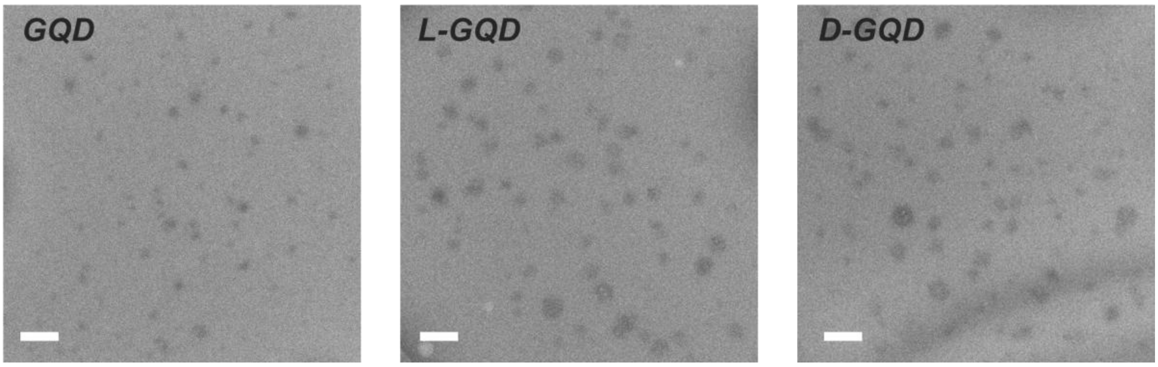


**Figure S7.** TEM image of **GQD**, ***L*-GQD** and ***D*-GQD** (100 μM) titrated with ICG (2 μM) in water. Scale bar: 100 nm.

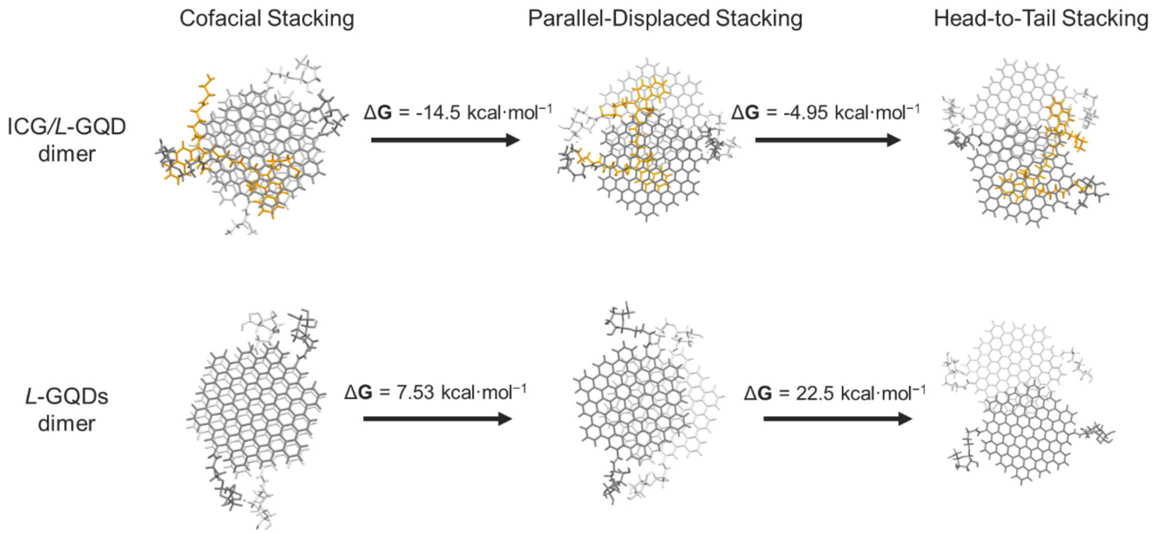


**Figure S8.** The Gibbs free energies (ΔG) of the optimized different stacked structures of *L*-GQD and ICG.

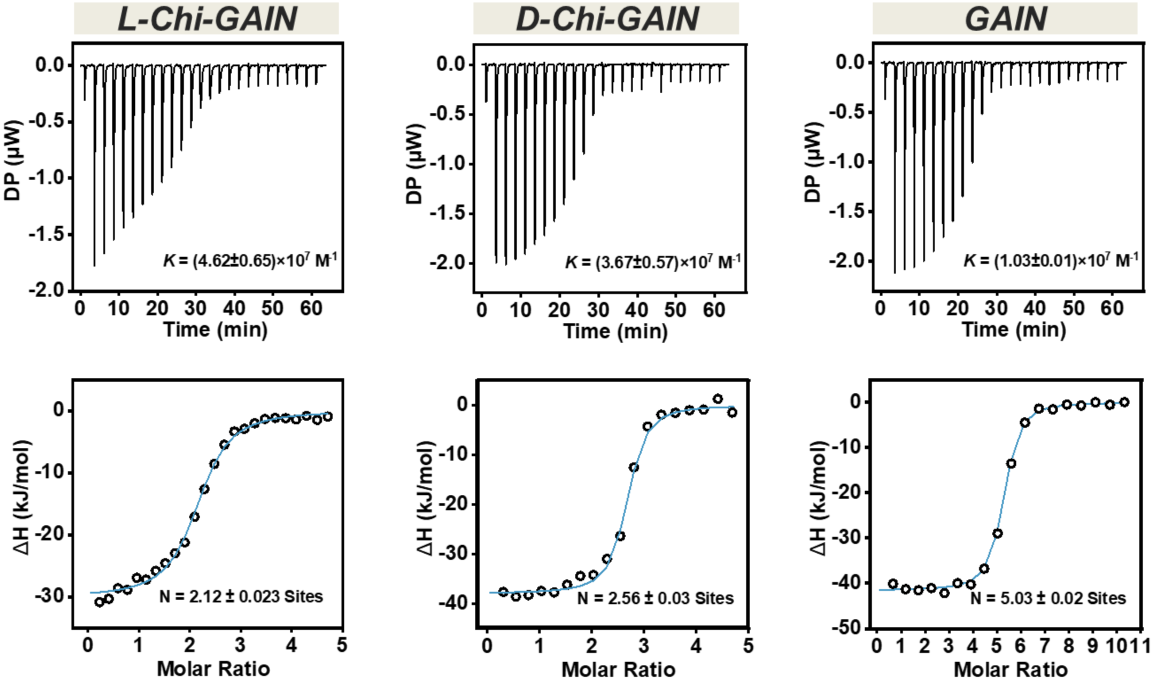


**Figure S9.** Isothermal Titration Calorimetry (ITC) of ***L*-GQD**, ***D*-GQD** and **GQD** (10 μM) titrated with ICG at 298 K in water.

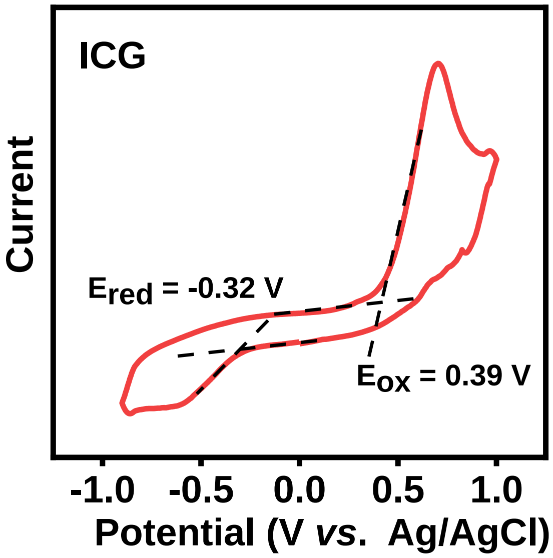


**Figure S10.** Cyclic voltammetry curves of ICG in water containing 0.1 M KCl, Ag/AgCl as the reference electrode, glassy-carbon electrode as the working electrode and Pt wire as the counter electrode; scan rate, 100 mV $s^{-1}$.

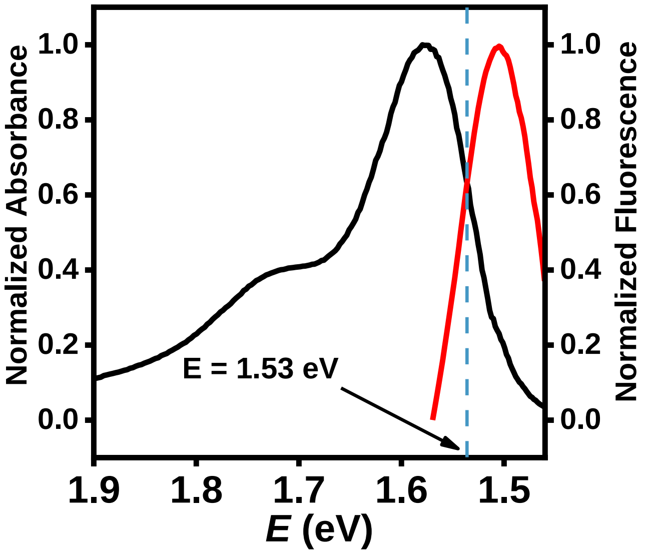


**Figure S11.** Normalized absorbance and emission spectra of ICG in water after converting the wavelength axis to an energy scale.

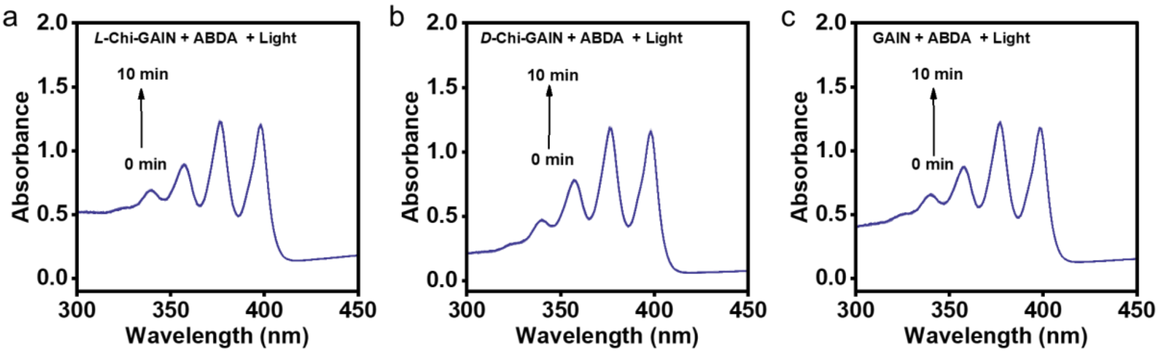


**Figure S12.** Absorption spectra of 9,10-anthracenediyl-bis(methylene)dimalonic acid (ABDA) (20 μM) after irradiation for different times in the presence of (a) ***L*-Chi-GAIN**, (b) ***D*-Chi-GAIN** and (c) **GAIN**.

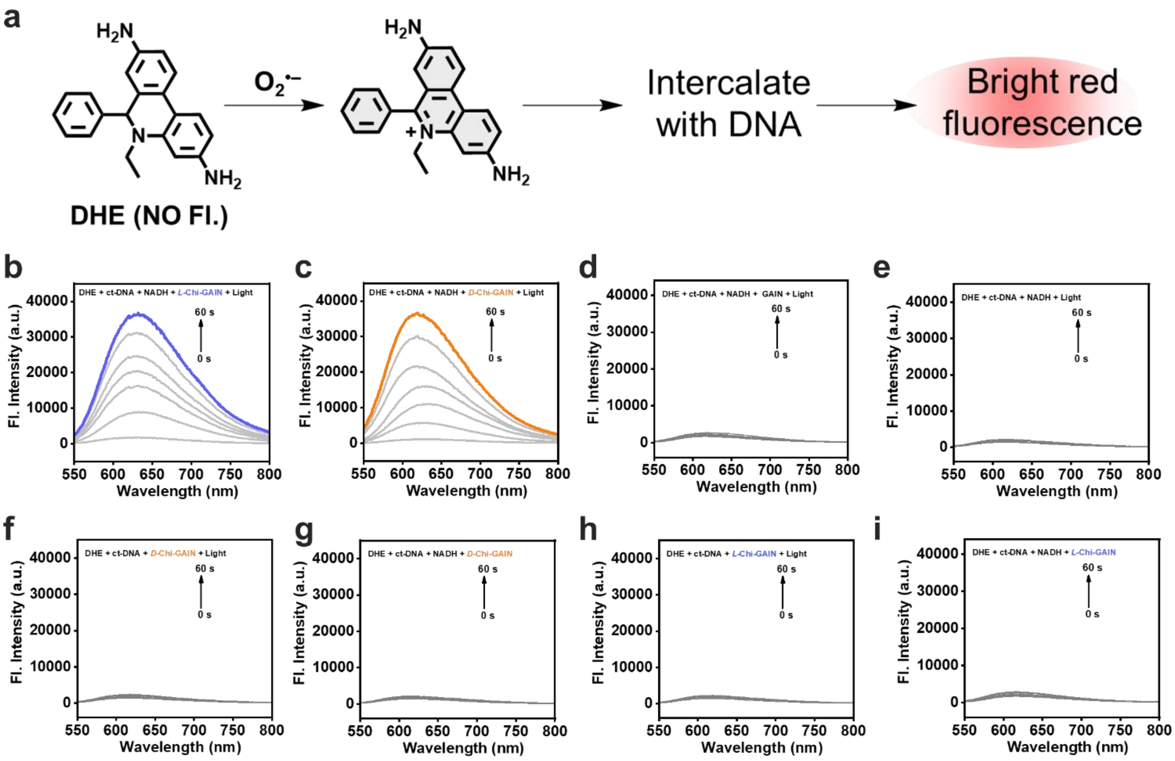


**Figure S13.** (a) The mechanism of Dihydroethidium (DHE) detects $O_2^{\cdot-}$ in the solution. Fluorescence spectra of DHE (50 μM) containing 500 μg/mL ctDNA at different conditions. (b) NADH + ***L*-Chi-GAIN** + Light; (c) NADH + ***D*-Chi-GAIN** + Light; (d) NADH + **GAIN** + Light; (e) NADH + light; (f) ***D*-Chi-GAIN** + Light; (g) NADH + ***D*-Chi-GAIN**; (h) ***L*-Chi-GAIN** + Light (i) NADH + ***L*-Chi-GAIN**.

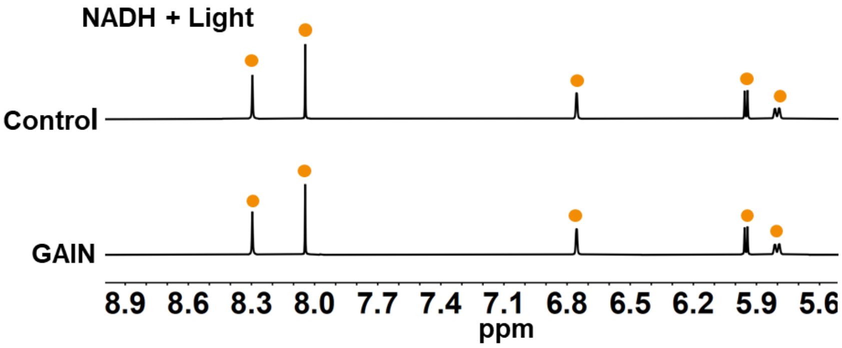


**Figure S14.** $^{1}H$ Nuclear Magnetic Resonance (NMR) of NADH with **GAIN** in $D_2O$ after irradiation.

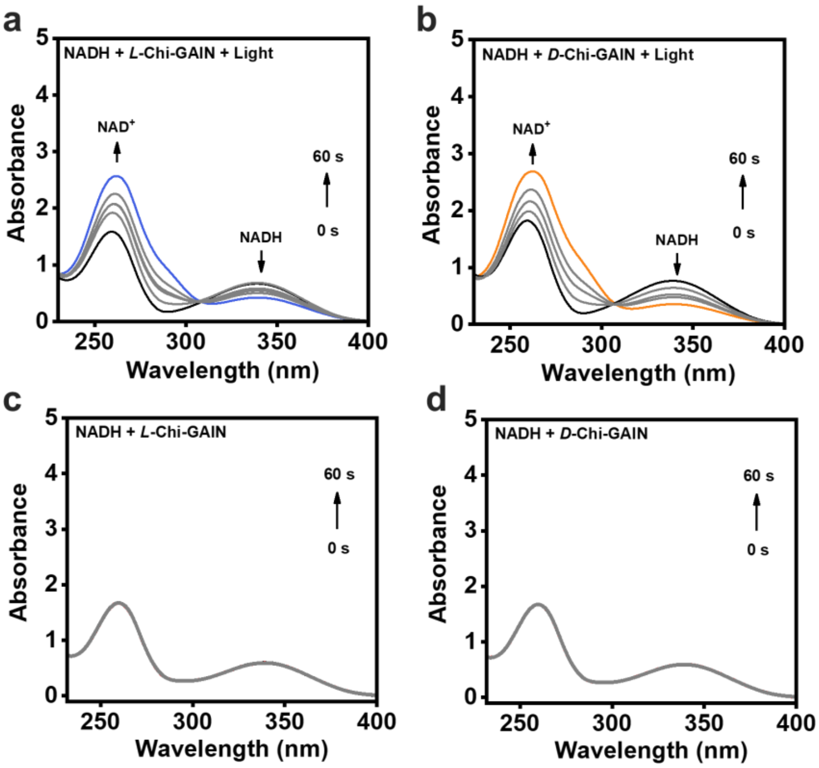


**Figure S15.** The absorption spectra of NADH (100 μM) under different conditions.

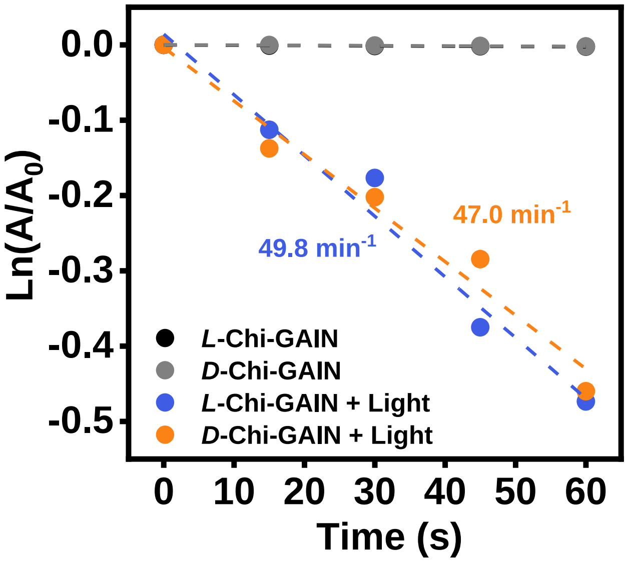


**Figure S16.** Plots of lnA/A₀ of NADH at 339 nm for different time intervals.

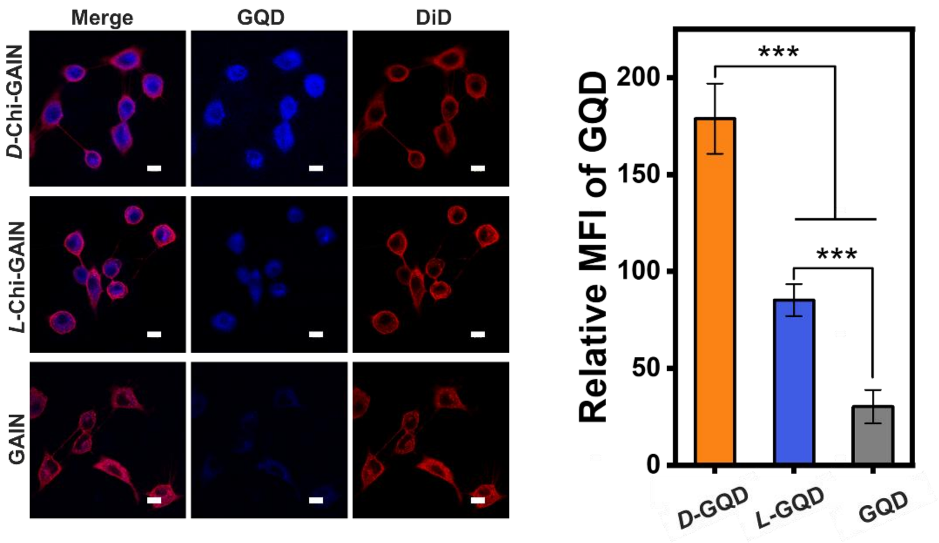


**Figure S17.** Confocal laser scanning microscopy (CLSM) imaging and GQD fluorescence quantification of HepG2 cells after treatment with ***D*-Chi-GAIN**, ***L*-Chi-GAIN** and **GAIN** (5 μM) for 4 h. DiD was used as a cell membrane marker. Scale bars: 10 μm. (*n* = 5 ), Data are expressed as the mean±SD. Statistical differences were analyzed by a Student's two-sided t-test. ***$P$<0.001.

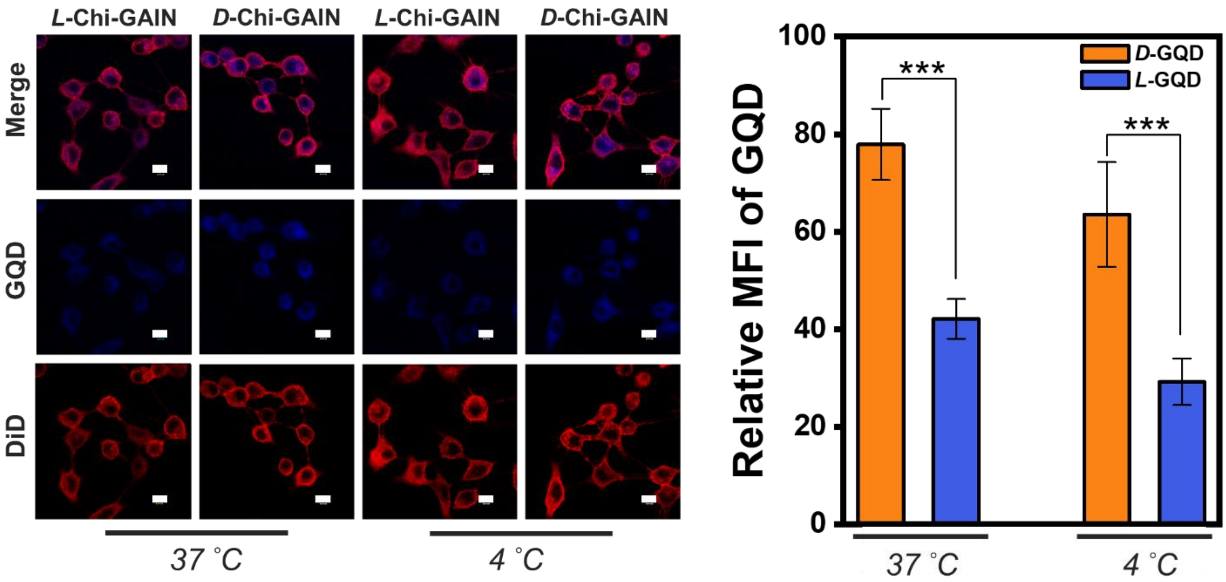


**Figure S18.** CLSM imaging and GQD fluorescence quantification of HepG2 cells pre-blocked with an ASGPR antibody and subsequently incubated with ***D*-Chi-GAIN**, ***L*-Chi-GAIN** (5 μM) for 4 h at 37 °C or 4 °C. DiD was used as a membrane marker. Scale bars: 10 μm.. ($n$ = 5 ), Data are expressed as the mean±SD. Statistical differences were analyzed by a Student's two-sided t-test. ***$P$<0.001.

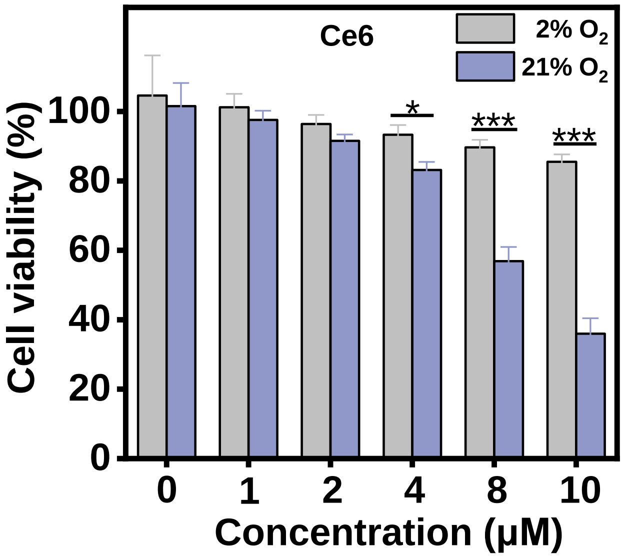


**Figure S19.** Cell viability of HepG2 cells subjected to the arrangement of Ce6 concentration in the presence of light irradiation under normoxia (21% $O_2$) or hypoxia (2% $O_2$). ($n$ = 5, mean ± SD) *$P$ < 0.05, ***$P$ < 0.001 determined by unpaired two-sided student $t$-test.

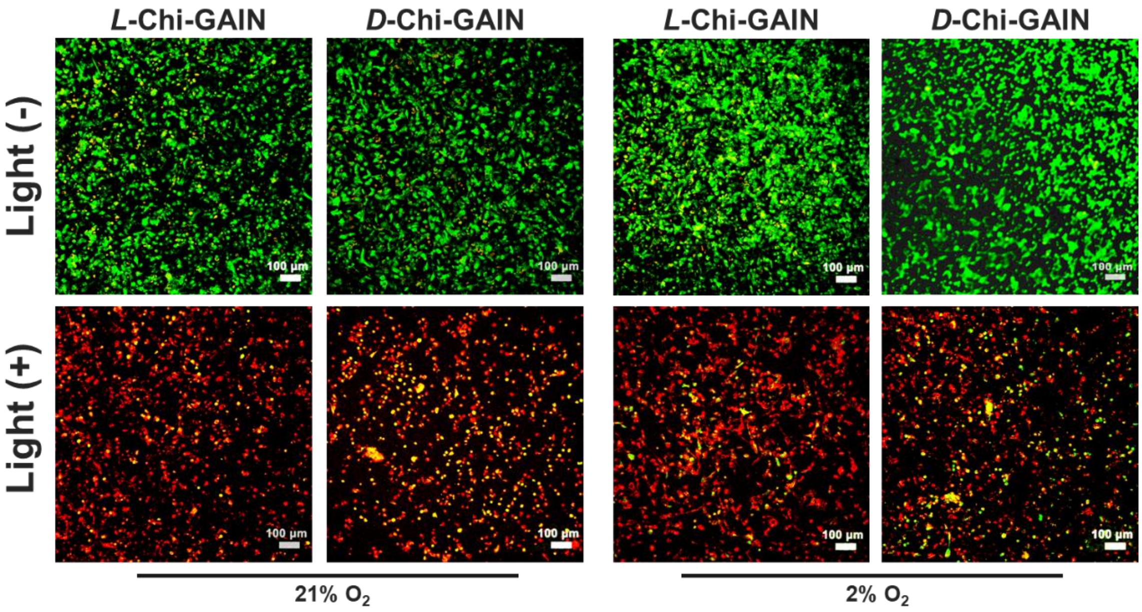


**Figure S20.** CLSM images of HepG2 cells treated with **Chi-GAIN (**20 µM**)** under normoxic/hypoxic conditions with/without light irradiation. Cells were stained with calcein-AM to identify live cells and with propidium iodide (PI) to identify dead cells. Scale bars: 100 µm.

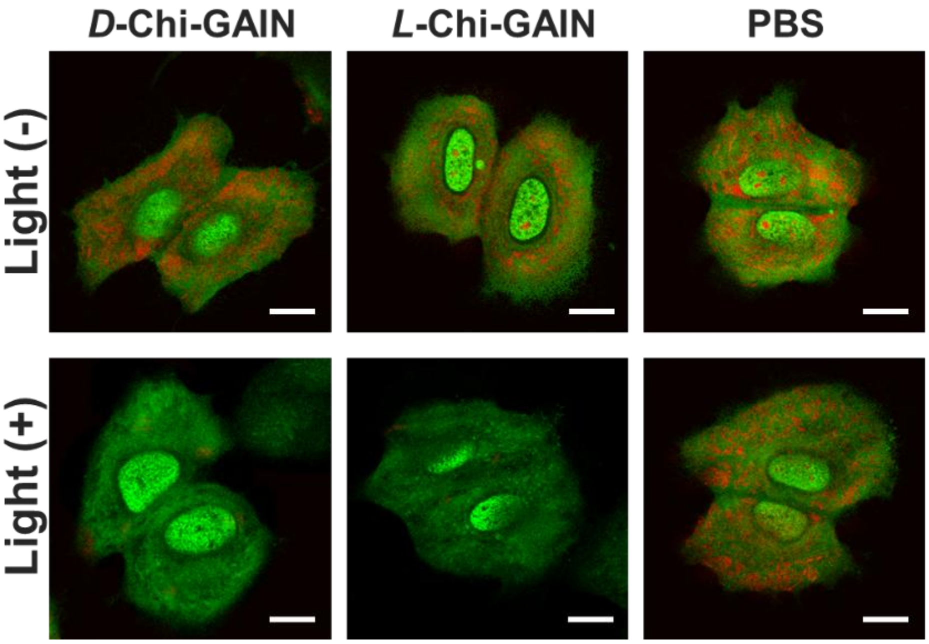


**Figure S21.** CLSM images of acridine orange (AO) staining for lysosomal integrity. Scale bars: 10 µm.

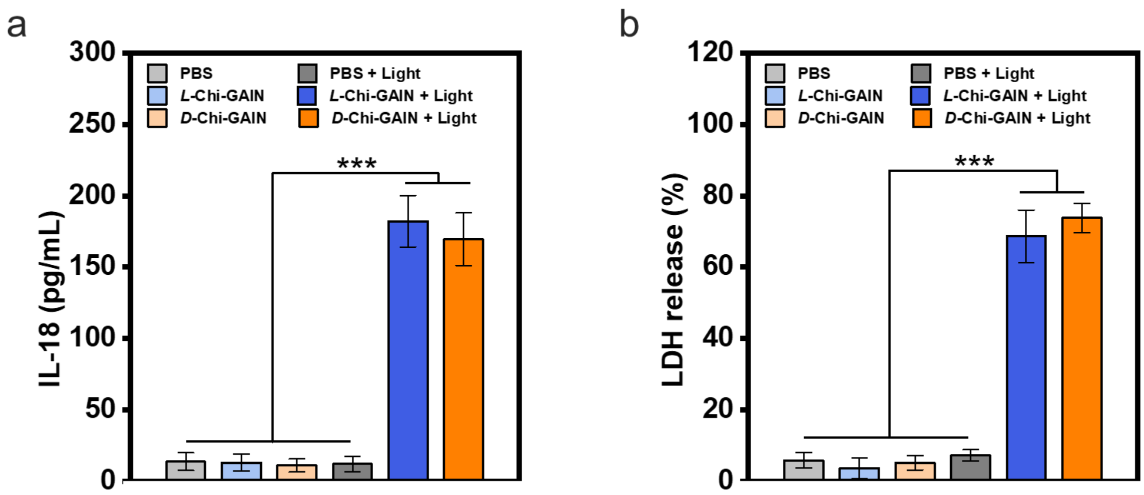


**Figure S22.** Quantification of (a) IL-18 and (b) lactate dehydrogenase (LDH) release from HepG2 cells after different treatments. Data are shown as mean ± s.d. (n = 5). Statistical differences were analyzed by two-tailed Student's t-test.

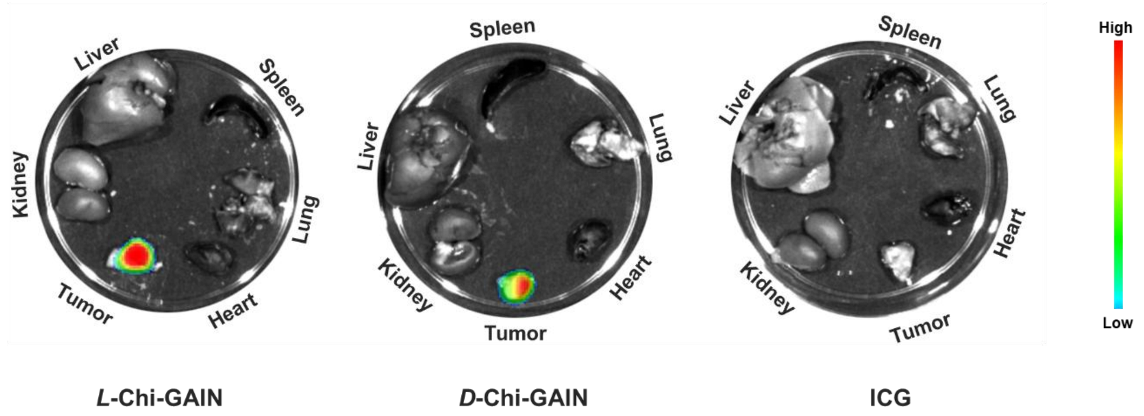


**Figure S23.** *Ex vivo* fluorescence imaging of the tumor, heart, liver, spleen, lung, and kidney after injection of ***L*-Chi-GAIN,** ***L*-Chi-GAIN** or **ICG** for 24 hours.

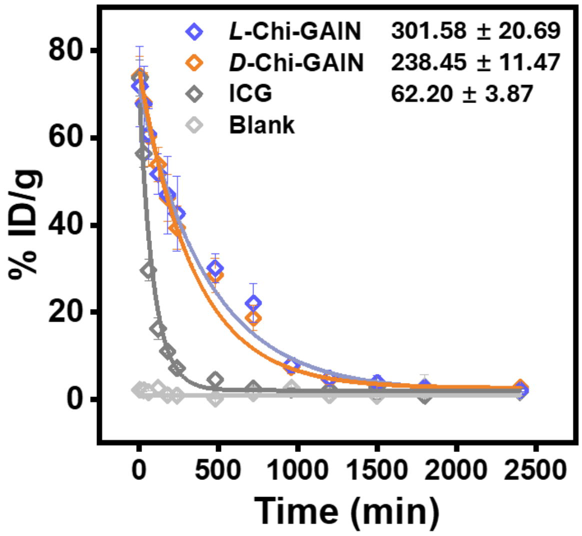


**Figure S24.** Concentration of ***L*-Chi-GAIN, *D*-Chi-GAIN** or **ICG** in plasma (% ID g$^{-1}$) determined by fluorescence measurements after injection ($n$ = 5, mean ± SD).

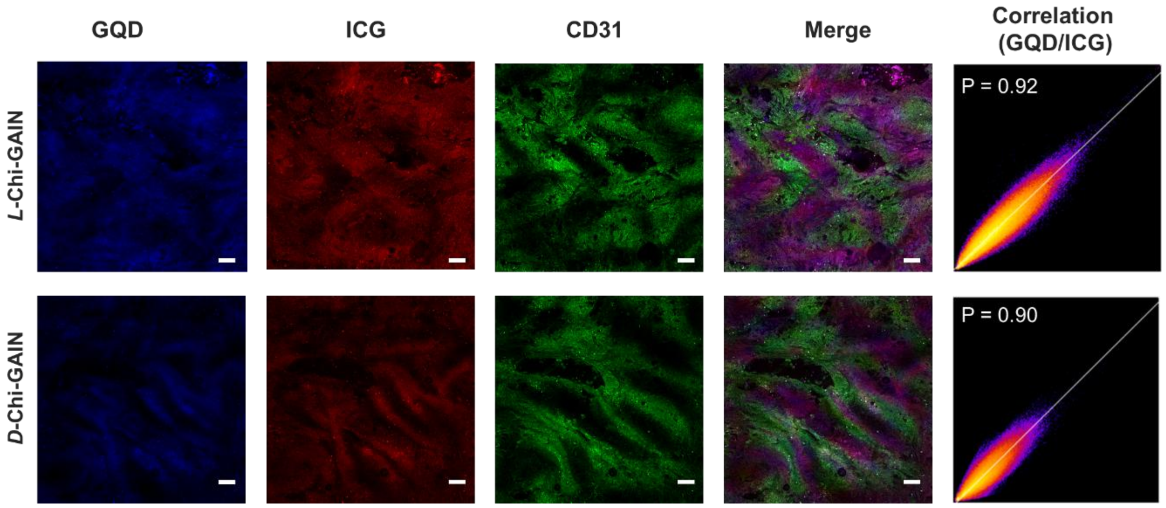


**Figure S25.** Fluorescence images of penetration of ***L*-Chi-GAIN** and ***D*-Chi-GAIN** into deep tumor tissues. Blue, GQD; green, blood vessels; red, ICG. Scale bar: 100 μm.

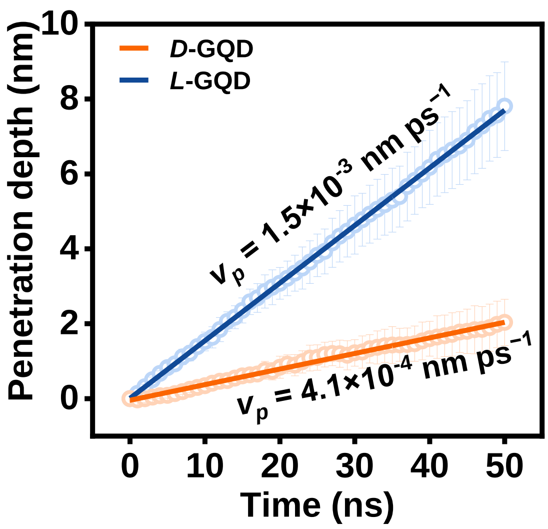


**Figure S26.** The penetration depth of *L*- and *D*-GQDs in ECM model at different times (mean ± SD, n = 5).

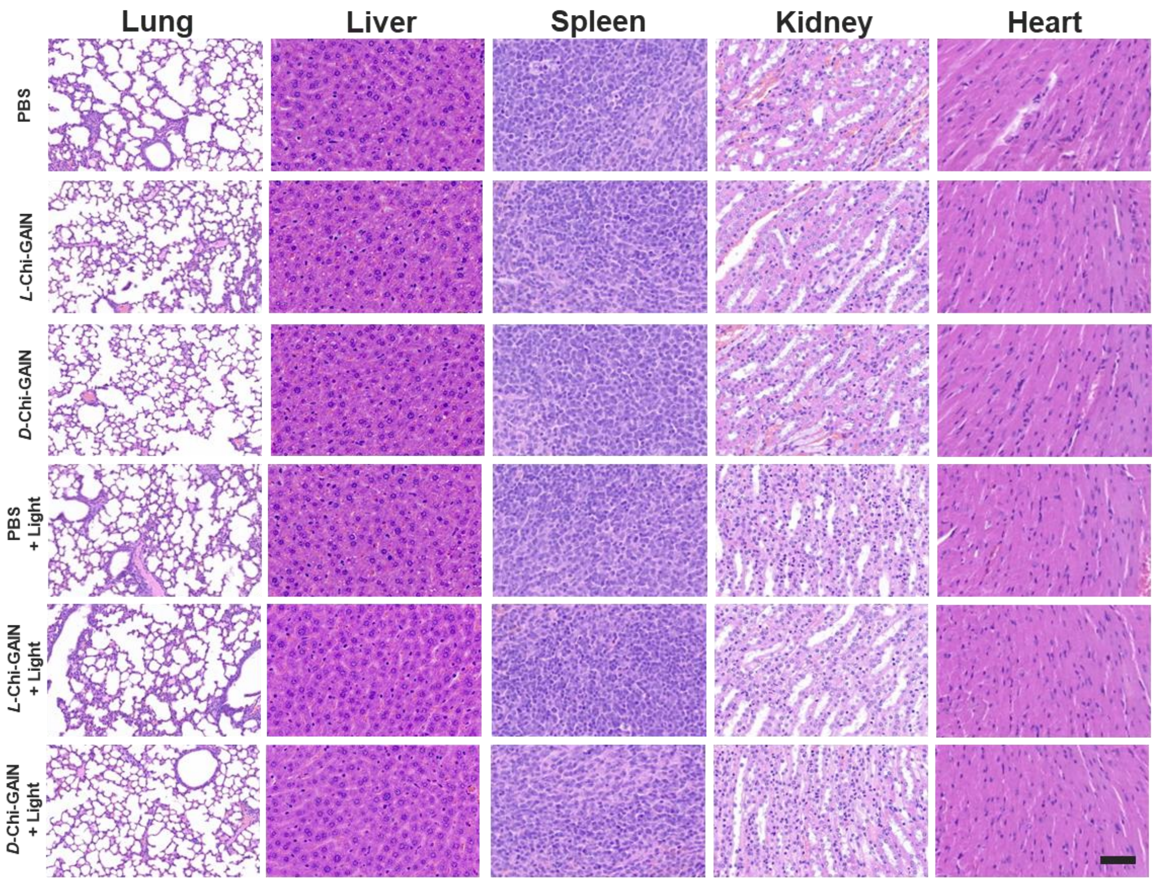


**Figure S27.** Hematoxylin and eosin (H&E) staining of main mouse organs (lung, liver, spleen, kidney, heart) slides from mice after different treatments. Scale bars: 50 μm.
